# Evolution of the cytochrome-*bd* type oxygen reductase superfamily and the function of cydAA’ in Archaea

**DOI:** 10.1101/2021.01.16.426971

**Authors:** Ranjani Murali, Robert B. Gennis, James Hemp

## Abstract

Cytochrome *bd*-type oxygen reductases (cytbd) belong to one of three enzyme superfamilies that catalyze oxygen reduction to water. They are widely distributed in Bacteria and Archaea, but the full extent of their biochemical diversity is unknown. Here we used phylogenomics to identify 3 families and several subfamilies within the cytbd superfamily. The core architecture shared by all members of the superfamily consists of four transmembrane helices that bind two active site hemes, which are responsible for oxygen reduction. While previously characterized cytochrome *bd*-type oxygen reductases use quinol as an electron donor to reduce oxygen, sequence analysis shows that only one of the identified families has a conserved quinol binding site. The other families are missing this feature, suggesting that they use an alternative electron donor. Multiple gene duplication events were identified within the superfamily, resulting in significant evolutionary and structural diversity. The CydAA’ cytbd, found exclusively in Archaea, is formed by the co-association of two superfamily paralogs. We heterologously expressed CydAA’ from *Caldivirga maquilingensis* and demonstrated that it performs oxygen reduction with quinol as an electron donor. Strikingly, CydAA’ is the first isoform of cytbd containing only *b*-type hemes shown to be active when isolated, demonstrating that oxygen reductase activity in this superfamily is not dependent on heme *d*.

## Introduction

The predominance of oxygen in our atmosphere determines the bioenergetic importance of oxygen as an electron acceptor and the prevalence of aerobic respiratory chains. There are only three enzyme superfamilies capable of acting as terminal respiratory oxygen reductases - heme-copper oxygen reductases, alternative oxidases and cytochrome *bd*-type oxygen reductases (cytbd)^1^. While enzymes from this superfamily have been characterized from a number of Bacteria, their role in archaeal respiration has not yet been determined. Archaeal aerobic respiratory chains share some similarities with bacterial respiratory chains, however they often differ in their composition of respiratory enzymes and are adapted to use different cofactors such as methanophenazine and F_420_. Complexes that are typically involved in bacterial respiration such as succinate-quinone oxidoreductases^2^, cytochrome *bc_1_* complexes^3^ and heme-copper oxygen reductases from archaea have been previously characterized^4,5^, while NADH:quinone oxidoreductases and alternative complex III are absent from this domain^6^. The presence of cytochrome *bd*-type oxygen reductases has been noted in archaeal genomes^7^, metagenomes and metaproteomes^8–11^ but, no functional member of the cytbd superfamily in archaea has ever been demonstrated.

Cytochrome *bd*-type oxygen reductase is a respiratory enzyme that converts oxygen to water using three hemes, unlike the heme-copper oxygen reductases which have two hemes and a copper in the active sites^1^. Purified cytbd accepts electrons from quinols using a low-spin heme *b*_558_ and transfers these electrons to a di-heme active site containing two high-spin hemes. In some of the characterized cytochrome *bd* enzymes, these active site hemes were shown to be heme *b_595_* and heme *d*, but some other isoforms were shown to contain only hemes *b*. Those cytochrome *bd* family members that contain only hemes *b* are usually referred to as cyanide insensitive oxidases (CIO) or cytochrome *bb*’–type oxygen reductase, and have been identified in *Pseudomonas aeruginosa, Bacillus subtilis* and others^12–15^. It is unclear whether the presence of only hemes *b* has a physiological implication but it has been suggested that these enzymes are less sensitive to inhibition by cyanide^13^. There is no sequence signature that distinguishes those enzymes in the superfamily that only contain heme *b*. No CIO has ever been isolated and characterized.

The canonical cytochrome *bd* oxygen reductases contain a minimum of two subunits, cydA and cydB, but often contain additional “auxiliary” subunits ^16–18^ such as CydX, a single-transmembrane subunit that is associated with cytochrome bd-I from *E. coli* that has been implicated in the stability of the enzyme^19^. Cytochrome *bd*-type oxygen reductases have a high affinity for oxygen^20^ and the previously characterized cytbds have been associated with roles in oxygen detoxification, respiratory protection of nitrogenases and as part of sulfide oxidizing respiratory chains^21–25^. Cytochrome *bd* catalytic turnover generates a proton motive force by translocation of protons using a conserved proton channel from the cytoplasm to the site of oxygen reduction located near the periplasmic side (electrically positive) of the membrane^26^. Yet, cytochrome *bd* is not as energetically efficient as the heme-copper oxygen reductases which pump protons in addition to translocating “chemical” protons from the cytoplasm to the periplasmic active site^27,28^. Expression of cytochrome *bd* has often been associated with microoxic conditions where a high-affinity oxygen reductase would be required^29^.

In this work, we used phylogenomics to determine the diversity and distribution of this high affinity oxygen reductase in Archaea and Bacteria. We determined that there are three distinct families of cytbd – one of which contained the quinol binding characteristics present in the structures of cytbd from *Escherichia coli* and *Geobacillus thermodenitrificans^30–32^* and two which do not – and discussed their evolutionary relationships. The distribution of these families even within Archaea involve significant variation and include the two distinct isoforms CydAB and CydAA’, the latter of which appears to have been created by gene duplication. We evaluate the relative distribution of the CydAA’ and CydAB within the domain archaea and consider the likely role of CydAA’ variants within their ecological context. In addition, we show that the CydAA’ from *Caldivirga maquilingensis* is a highly active oxygen reductase with unique biochemical and structural characteristics. This combined phylogenomic and experimental approach has significantly expanded our knowledge of the evolutionary and biochemical diversity within the superfamily, which has important implications for the role of the cytbd superfamily in novel respiratory pathways.

## Results

### Diversity of cytochrome *bd*-type oxygen reductases

The molecular structures of cytochrome *bd*-type oxygen reductases from *Escherichia coli* and *Geobacillus thermodenitrificans* have been determined, and showed that cytbd typically has two conserved subunits cydA and cydB, along with a third subunit, cydX or cydS which is a single transmembrane subunit that is not well conserved or found along with cydA and cydB in the genome^30–33^. Of the two main subunits, cydA is better conserved in all known cytochrome *bd*-type oxygen reductases while cydB is very divergent and is hypothesized to have evolved at faster rates than cydA^34^. The cydA subunit is made of nine transmembrane helices and contains almost all the conserved amino acids known to be important for catalyzing oxygen reduction and proton translocation, including the ligands for three hemes and the residues forming a proton channel^35,36^. The first four helices of cydA contain all of the amino acids that form the active site. These include the proton channel and ligands to bind the active site heme *b_595_* and heme *d* (these ligands have only been verified in the isoforms containing hemes *b* and *d*, and not the ones containing only hemes b). The other five helices (V-IX) form the quinol binding site in the biochemically characterized *bd*-type oxygen reductases and include the ligands to heme *b_558_*, the point of entry for electrons from quinols^30–32^. With this structural framework in mind, we performed a sequence analysis of cytochrome *bd*-type oxygen.

An analysis of 24706 genomes available in the Genome Taxonomy Database (release89)^37,38^, revealed the presence of 17852 cydA homologs. Of these, 13007 genomes contained at least one cydA homolog, suggesting that this enzyme family is widely distributed and important (**Supplementary Table1**). Phylogenomic analysis of cydA homologs revealed 15 clades of cydA that could be distinguished on the basis of unique sequence characteristics.^1^ (**Supplementary Figure S1**, **Supplementary Tables1,2**). Four of these clades contain the features that are considered part of the quinol binding site – for e.g., conserved residues Lys252, Glu257 (*E.coli* cydA numbering) while the remaining do not. We inferred that the former four cydA clades were quinol:O_2_ oxidoreductases and we named them qOR1, qOR2, qOR3 and qOR4a. While cydA of the families qOR1, qOR2 and qOR3 associate with cydB to form cydAB, cydAA’ is formed by the co-association of two distinct cydA clades, qOR4a and qOR4b. qOR4b does not possess quinol binding site characteristics and is likely the result of a gene duplication event. Phylogenetic clustering of cydA sequences from all 15 cydA clades demonstrated that 2 of the remaining clades are missing quinol binding sites and instead contain a number of heme *c* binding motifs (CxxCH) (**Supplementary figure S1, Supplementary multiple sequence alignments MSA1, MSA3**). We named these enzymes OR-C1a and OR-C1b because of the presence of heme *c* binding motifs. Similar to qOR4a/4b, the ‘a’ and ‘b’ attachment to the names signifies that their genomic context suggests that they co-associate to form one enzyme OR-C1. Eight of the remaining cydA clades were related and named OR-N1, OR-N2, OR-N3a, OR-N3b, OR-N4a, OR-N4b, OR-N5a and OR-N5b. OR-N is named for Nitrospirota because of predominance of these enzymes in that phylum (**Supplementary Figure S1, Supplementary multiple sequence alignments MSA1, MSA2, MSA3**). Their close relationship is also supported by the likely structure of the proteins of which they are a part and their genomic context (**Figure 1**). We have attempted to develop a nomenclature for the cytochrome *bd*-type oxygen reductase family that can be easily expanded upon. We designate 3 large families of cytochrome *bd*-type oxygen reductases – qOR, OR-C and OR-N - based on their phylogenetic placement, presence/absence of biochemical signatures such as quinol and heme *c* binding site features, genomic operon context and taxonomic origin. We have designated subfamilies numerically starting from 1 and attached an ‘a’ or ‘b’ subscript if it is likely that two cydA subfamilies co-associate to form one enzyme. Most of the ‘a’-type subfamilies include the proton channel residues E99 and E107 (*E. coli* numbering) while ‘b’-type subfamilies do not. It appears that ‘a’ and ‘b’-type subfamilies are also the result of multiple independent gene duplication events within this superfamily. We will discuss the unusual number of gene duplication events within the cytbd superfamily and the OR-C and OR-N families later in the text and in **Supplementary Material** but begin with the quinol oxidizing qOR family. Sequences from this family contain all the amino acids that were previously identified as forming the heme ligands, proton channel, oxygen reduction site and quinol binding site^30–32,36^.

**Figure 1.**
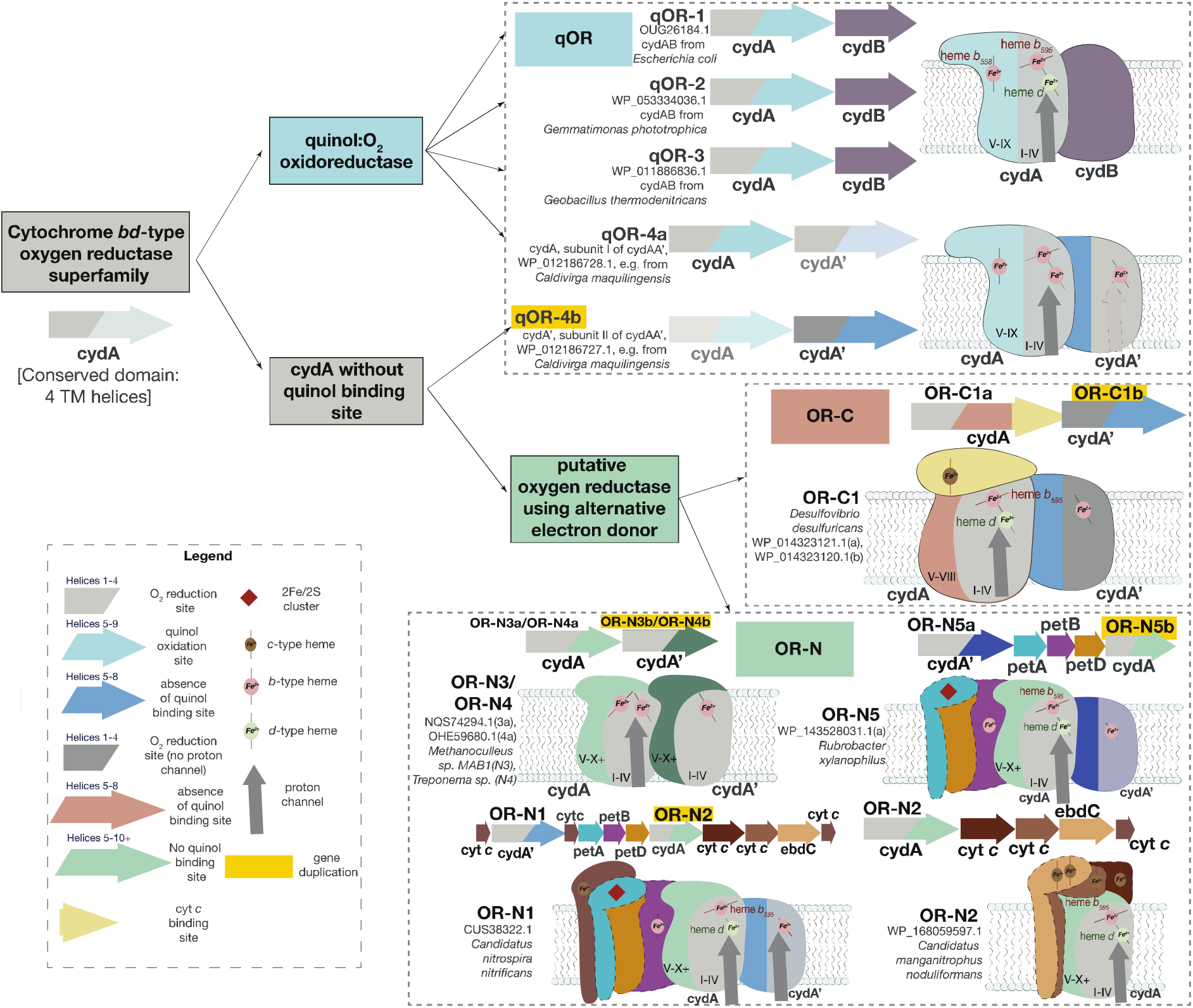
Diversity of the cytochrome *bd* oxygen reductase superfamily. The cytochrome *bd* oxygen reductase superfamily is divided into 3 families based on phylogenetics and structure – qOR, OR-C and OR-N. qOR is defined by the presence of the quinol binding site in subunit I (cydA). OR-C is missing the quinol binding site but has a heme *c* binding site in subunit I. OR-N is also missing the quinol binding site and is commonly found in operons containing alternative electron donors. Various subfamilies within each family are also shown (Supplementary Figure 1). The operon context and putative protein complex arrangement of each cydA-containing enzyme is also shown with a reference protein accession number and source microorganism. The potential gene duplication events are highlighted in yellow. A legend is also provided to mark the related conserved domains in the same colors and redox co-factors such hemes and iron-sulfur clusters. A more detailed explanation of the figure including a description of the various subunits and characteristics of the families and subfamilies is provided in **Supplementary Material**.

### Evolution of the quinol-oxidizing cytochrome *bd*-type oxygen reductases (qOR)

To explore the evolutionary relationship between the families qOR1, qOR2, qOR3 and qOR4a, which are true orthologs, we generated a maximum likelihood phylogenetic tree using RAxML with the OR-C and OR-N family sequences as outgroup (**Figure 2**). Sequence features can be identified to distinguish these families and to validate the above identified monophyletic clades as meaningfully distinct; some of which are outlined below while the remaining features are mentioned in **Supplementary Table 6**. All cydA sequences from qOR1 subfamily have 7 amino acids between the two conserved glutamates in the proton channel Glu99 and Glu107 such as in *Escherichia coli* cydA ^26,39^ while cydA sequences from qOR2, qOR3 and qOR4a typically have 6 amino acids between the two conserved glutamates (ex. as between Glu101 and Glu108 in qOR3-subfamily cytbd from *Geobacillus thermodenitrificans^30^*). This insertion/deletion has been hypothesized to lead to a reversal in the position of hemes from the qOR1-bd in *Escherichia coli* to the qOR3-bd in *Geobacillus thermodenitrificans* although further research is required to establish whether the reversal of heme positions is universal (further discussion on the insertion/deletion in the proton channel and the Q-loop is included in **Supplementary Material**). Sequence features which distinguish the qOR4a-subfamily cytbd are insertions between helices V and VI, as well as insertions in helix VIII (**Supplementary alignment MSA1, Supplementary Table 6**) Conserved tyrosines (Tyr115 and Tyr117 *Geobacillus thermodenitrificans* cydA numbering) are present in qOR2 and qOR3 families but not in the qOR4a-subfamily, which is consistent with the close evolutionary relationship between the qOR2 and qOR3-subfamilies observed in the tree topology. Other conserved sequence features, unique to each family are listed in **Supplementary alignment MSA2** and **Supplementary Table 6**.

**Figure 2.**
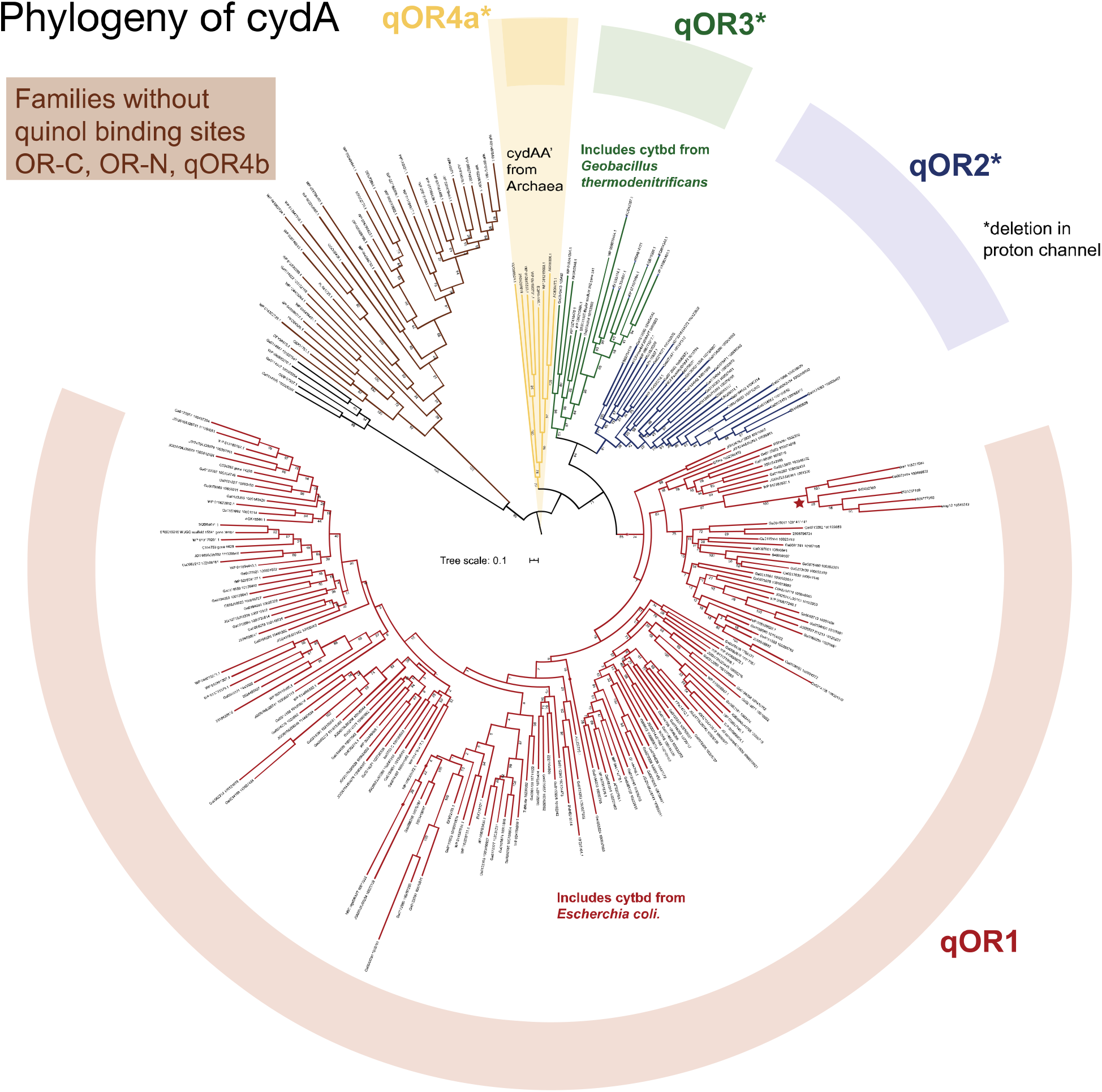
Phylogeny of quinol-oxidizing cytochrome *bd*-type oxygen reductases. At least four clades of quinol-oxidizing cytochrome *bd*-type oxygen reductases could be identified - qOR1, qOR2, qOR-3 and qOR-4a. The long branch within the qOR1 clade (red star) is comprised of sequences missing the proton channel that is conserved in all other quinol-oxidizing cytochrome *bd*-type oxygen reductases. Subunit I of cydAA’ is from the qOR4a family. The cytochrome *bd*- type oxygen reductases that do not contain the quinol binding site (OR-C, OR-N, and qOR-4b families) were used as the outgroup.

Comparing the cydA phylogenetic tree and the distribution of cytochrome *bd*-type oxygen reductases across Archaea and Bacteria provides some insight into the relative age of these families. The qOR1 subfamily, which includes the *Escherichia coli* enzyme, at present count seems to be the most widely distributed with enzymes in over 60 bacterial phyla, (**Supplementary Table 2, 4**) but it is only sparsely distributed in Archaea. In fact, there are only a very few representatives in Euryarchaeota and Asgardarchaeota (**Figure 2**). It is only widely distributed in Halobacterota, whose oxidative metabolism is expected to have evolved relatively late^40^(**Figure 3**). This strongly suggests that the qOR1 subfamily is the oldest of the extant families and that it is likely that cytochrome *bd*-type oxygen reductases originated in Bacteria. While cydA from the qOR2 subfamily is also fairly well-distributed and found in over 20 bacterial phyla, the qOR3-subfamily enzymes are almost exclusive to the Firmicutes and Firmicutes_I phyla with a few enzymes in Archaea. The qOR4a-subfamily enzymes appear to be specific to the Archaea (**Supplementary Tables 2, 4**). A close evolutionary relationship between the qOR2 and qOR3-subfamilies is suggested by cydA tree topology and identifiable sequence characteristics but other trees we inferred have modelled a closer relationship between the qOR2 and qOR1 subfamilies (data not shown). Furthermore, the qOR1 subfamily has 7 amino acids between the conserved glutamates in the proton channel, while the qOR4a, qOR2 and qOR3 subfamilies consistently have 6 amino acids. Lastly, enzymes from the qOR4a, qOR2 and qOR3 subfamilies are almost completely absent from Proteobacteria. This suggests that the qOR2, qOR3 and qOR4a subfamilies diverged from the qOR1 family, Before proteobacteria diverged from other bacterial phyla. While our dataset and phylogenetic analysis is consistent with the above discussion, it must be noted that many lateral gene transfers have been observed within the cytochrome *bd*-type oxygen reductase^1^ which complicate evolutionary analysis.

**Figure 3.**
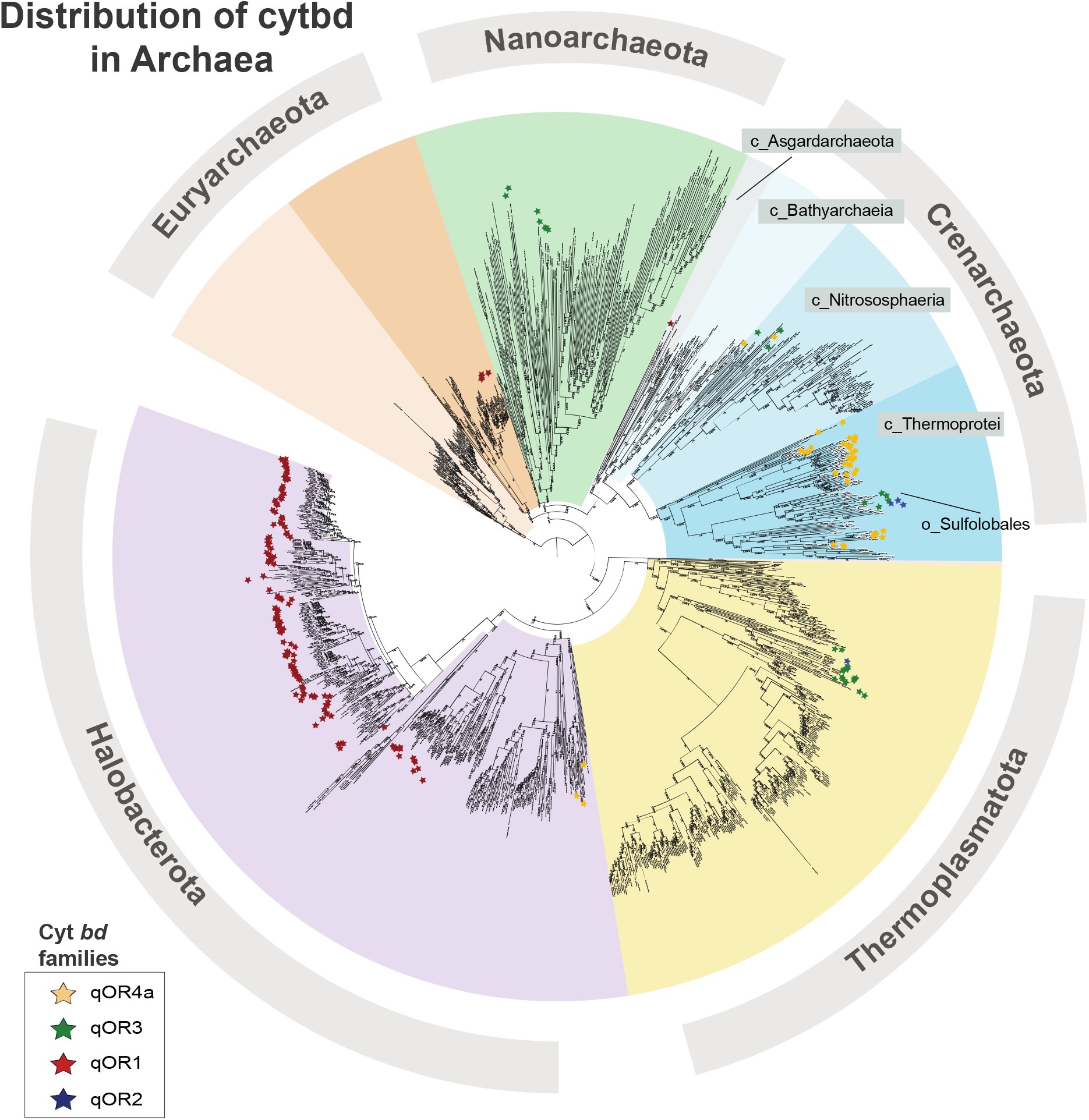
Distribution of cytochrome *bd*-type oxygen reductases in Archaea. Cytochrome bd-type oxygen reductases are sporatically distributed throughout the Archaea. The qOR4a family (cydAA’) is predominantly found within the *Thermoproteales* and *Desulfurococcales* orders of Crenarchaeota.

As mentioned above, most cydA subfamilies are widely distributed within Bacteria and Archaea, but the qOR4a-subfamily is unique in having sequences that belong only to Archaea. In addition, the qOR4a-subfamily is unique in having a completely different subunit II (cydA’), while the qOR1-, qOR2- and qOR3-subfamily members appear to have cydB homologs as their subunit II. cydB is either not homologous to cydA’ or is evolutionarily distant. The unique ancestry of the qOR4a-subfamily enzymes which is specific to archaea raises a question about its distribution within that domain.

### Distribution of cytochrome-bd type oxygen reductases in archaea

To investigate the distribution of cytochrome *bd*-type oxygen reductases within Archaea and to contextualize the evolution of cydA within archaeal evolution, we mapped the presence of qOR4a, qOR1, qOR2 and qOR3 subfamilies of cydA onto a phylogenetic tree of all archaea, using a concatenated gene alignment made from the archaeal genomes in GTDB^37^ using Anvi’o^41^, (**Figure 2**). It is clear from this representation that most of the qOR4a-subfamily or cydAA’ belong to the class Thermoprotei within the phylum Crenarchaeota with a few cydAA’ in Nitrosphaeria, Thermoplasmatota and Archaeoglobi. Within the Thermoprotei, almost all members of the order Thermoproteales contain cydAA’ and family *Acidilobaceae* contain cydAA’ (**Supplementary Table 3**).

To place cydAA’ into an ecological context we looked at their environmental distribution (**Supplementary Table 5**). Microbes containing cydAA’ are largely found in solfataric fields, hot springs and deep-sea vents, suggesting that cydAA’ might only be utilized by thermophiles, such as organisms from the genus *Vulcanisaeta, Caldivirga, Thermofilum* and *Thermocladium^42^*. Within Yellowstone National Park (YNP), a number of these genera are found in hypoxic, sulfur/iron-rich ecosystems, although *Pyrobaculum* and *Thermofilum* have also been found in more oxygenated environments. It has been suggested that members of the Thermoproteales which are found in aerobic environments have a heme-copper oxygen reductase and are more likely to be using aerobic respiration as their primary energetic pathway^42^. This is consistent with what we observe in Thermoproteales – organisms which do not have cydAA’ have heme-copper oxygen reductases instead (**Supplementary Table 3**). However, of the 8 *Pyrobaculum* genomes in the GTDB database, the three genomes that have cydAA’, but are missing a heme-copper oxygen reductase are capable of aerobic respiration^43,44^. This is suggestive of an adaptation based on oxygen availability in the environment resulting in a trade-off between the greater energetic efficiency and higher oxygen affinity of HCOs and *bd* respectively^20,28^.

Expression of the cydAA’ genes have been demonstrated in the hot springs and sulfur-rich/iron-rich environments within Yellowstone National Park, using RT-PCR^10^ and metatranscriptomics (**Table 1**). While the hot springs were typically hypoxic and sulfur-rich, the iron oxide mats had higher oxygen concentration at the surface and had <0.3 μM concentrations of O_2_ within 1 mm. Nitrosphaeria, Acidilobaceae, Thermoproteales and Thermoplasmatota expressed cydAA’ in these environments, however it is not clear whether these microorganisms were exposed to high O_2_ concentrations. In fact, the *Acidolobaceae* are expected to be found in the middle and bottom layers of this mat where O_2_ concentrations are lower^10,45^. All of the above observations are consistent with the presence of cydAA’ in microaerobic and hypoxic environments. The obvious question that needed to be addressed is whether cydAA’ actually functions as an oxygen reductase. This was accomplished by biochemically characterizing the CydAA’ from *C. maquilingensis*.

**Table 1.**
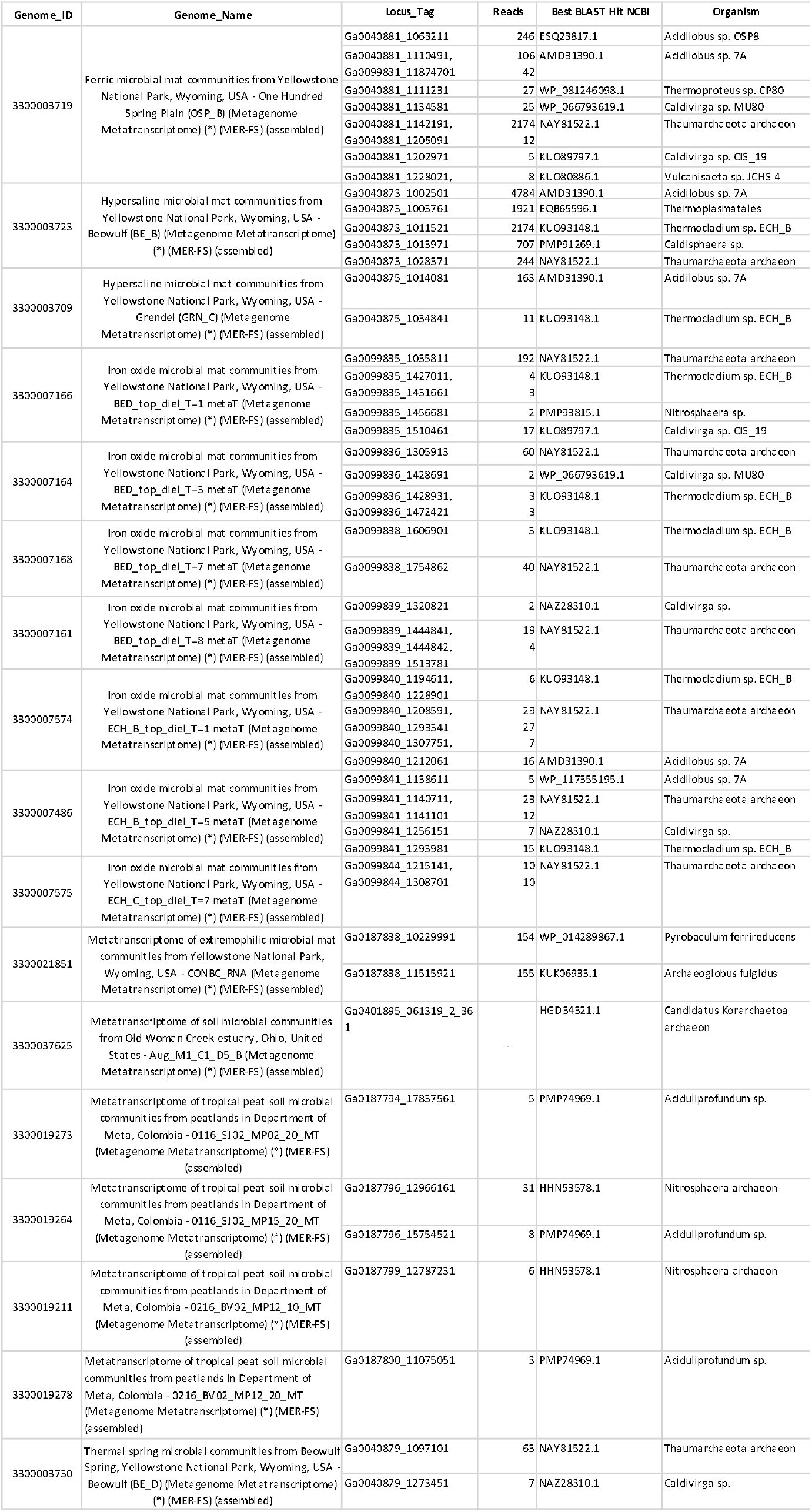
cydAA’ is expressed in many environments. Protein expression is estimated based on read counts in metatranscriptomes.

### Partial purification and spectroscopic characterization of the cydAA’ from *C. maquilingensis*

The cydAA’ operon from *Caldivirga maquilingensis* consists of two genes – *cydA* and *cydA*’. There are no additional subunits encoded within the operon corresponding to cydX/cydY or cydS, which are associated, respectively, with *E.coli* cydAB and *G.thermodenitrificans* cydAB. Homologues of these subunits are not apparent in the *C. maquilingensis* genome. We cloned the operon into the pET22b vector and expressed it in an *Escherichia coli* strain in which both bd-I and bd-II were deleted (CBO - C43, Δ*cyd*A Δ*appB*)^17^. The enzyme, cytochrome bb’ oxygen reductase from *Caldivirga* was engineered to have numerous different tags – 6xHistidine, FLAG, GST and GFP. None of these tags were successful, either because of a poor yield of protein or because of the inability of the affinity-tagged proteins to bind to columns with their corresponding epitopes. A GFP-tagged protein was used to verify the expression in *E.coli* of CydAA’ from *Caldivirga maquilingensis*. The presence of the protein could be observed by following the fluorescence of the protein under UV light. Since subunit II was tagged with GFP, it confirms the presence of subunit II in the preparation (**Supplementary Figure 3**). In addition, the purified protein was verified by mass spectrometry with many peptides recovered from subunit I. (**Supplementary Figure 4**). Gel electrophoresis of a partially purified CydAA’ shows two bands of the sizes expected for CydA and CydA’ (**Supplementary Figure 5**).

A UV-visible spectrum of CydAA’ in the reduced-minus-oxidized state reveals the absence of the heme *d* absorbance peak. The presence of heme *b*_595_ is also not apparent in the spectrum since the maxima at 595 nm and the Soret peak at 440 nm are also missing. This could indicate that heme *b_595_* is low spin in this preparation. Hemes were extracted from CydAA’ of *Caldivirga maquilingensis* as described previously^46^. Only *b*-type hemes are present in the enzyme (**Figure 4**). This was verified by analyzing the hemes in the protein by LC-MS (data not shown).

**Figure 4.**
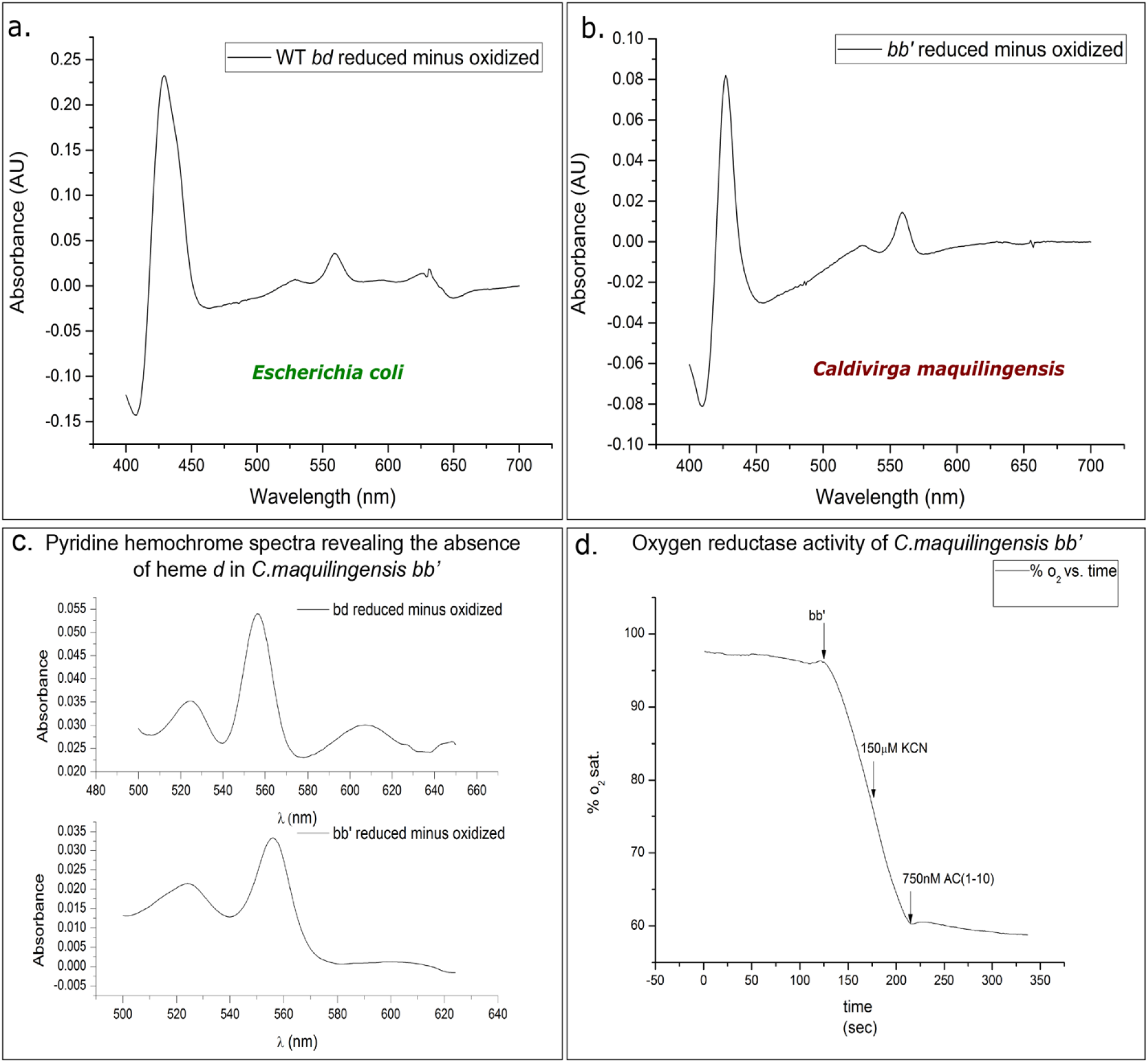
Biochemical characteristics of cytochrome bb’ (cytbb’) from *Caldivirga maquilingensis*. (A and B.) UV-visible spectra of cytochrome *bd*-type oxygen reductases purified from *Escherichia coli* and *Caldivirga maquilingensis*, respectively. C. Pyridine hemochrome spectra of *Caldivirga maquilingensis cytbb*’ reveals the absence of heme *d* in the partially purified enzyme. D. Oxygen reductase activity of cytbb’ from *C. maquilingensis* shows that it is highly active and cyanide insensitive. It is sensitive to Aurachin C1-10, a quinol binding site inhibitor which also inhibits *E. coli* cytochrome *bd*.

### CydAA’ from *Caldivirga maquilingensis* has oxygen reduction activity

The oxygen reduction activity of CydAA’ was tested using a Clark electrode, with reduced coenzyme Q1 (reduced using DTT) as the electron donor. (**Table 2**, **Figure 4**) At 37 °C the specific activity is ~330 e^-^/s (/heme *b*). While this is not as high as the activity of *E. coli bd* at the same temperature (over 1000 e^-^/s), the enzymatic activity is substantial, particularly considering the fact that the source of the enzyme is a thermophilic organism whose growth is optimum at 65 °C. The oxygen reductase activity of CydAA’ is insensitive to the presence of 250 μM KCN, a concentration of cyanide that would completely inhibit the activity of heme-copper oxygen reductases^1^. Since CydAA’ was expressed in the *bd*-deletion mutant, *E.coli* strain CBO, the only other potential oxygen reductase in this preparation is *bo_3_* ubiquinol oxygen reductase^47^ so the lack of cyanide sensitivity confirms that our purification protocol has separated the two enzymes. The enzyme is also susceptible to Aurachin AC1-10, a known inhibitor of cytochrome *bd* at concentrations as low as 250 nM^48^. We did not test for other possible functions for cydAA’ such as catalase activity^49^ or peroxidase activity^50^.

**Table 2.**
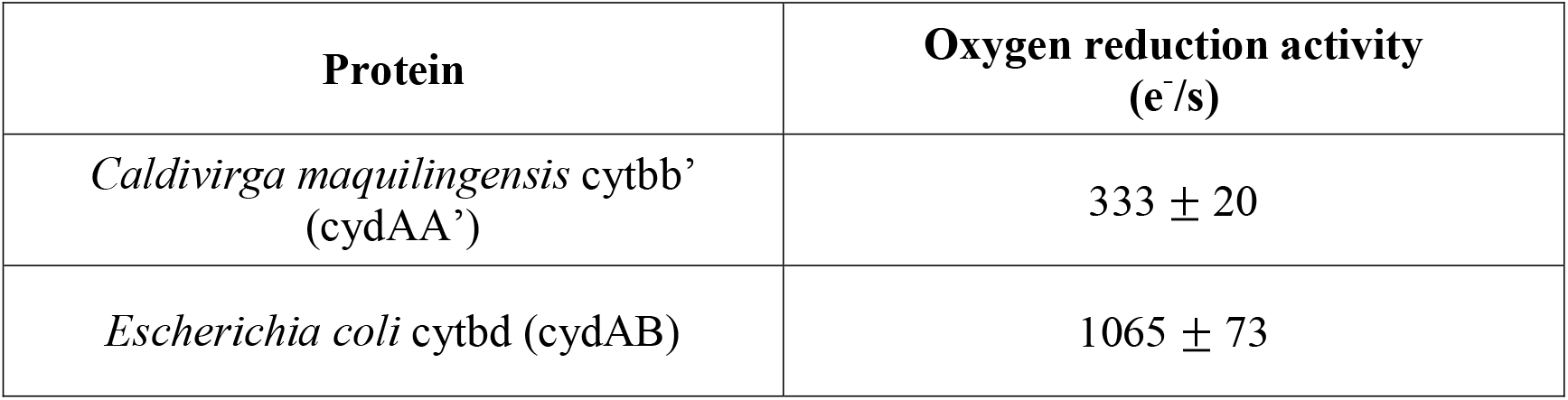
Oxygen reduction activity of *E. coli* cytbd and *C. maquilingensis* cytbb’ in the presence of 350 μM coenzyme Q1 and 5 mM DTT.

| **Protein** | **Oxygen reduction activity**<br>**(e<sup>-</sup>/s)** |
| --- | --- |
| *Caldivirga maquilingensis* cytb<sub>b</sub>'<br>(cydAA') | 333 ± 20 |
| *Escherichia coli* cytb<sub>d</sub> (cydAB) | 1065 ± 73 |

We previously noted that cydAA’ is typically found in organisms that perform sulfur-based chemistry such as sulfur reduction and sulfate reduction (**Supplementary Table 5**) and use DMSO-reductase like enzymes which use molybdopterin as a cofactor^10^. Combining the above observation with the demonstrated oxygen reductase activity of CydAA’, it is likely that the role of CydAA’ is to detoxify oxygen to protect oxygen-sensitive enzymes involved in sulfur metabolism. This is similar to its expected role in Desulfovibrio^24^ and in the protection of nitrogenases during the process of nitrogen fixation^22^.

It is striking that oxygen reduction is conserved in the qOR4a-subfamily despite the replacement of CydB with CydA’ and therefore, it is worth considering the similarities and differences between *E.coli* CydAB and *C.maquilingensis* CydAA’.

### Structural differences between CydAB and CydAA’ inferred from homology models

To aid in the understanding of differences between CydAA’ and other CydAB, we used multiple sequence alignments (**Supplementary Figure 2**) and structural models of cydA and cydA’ from *Caldivirga maquilingensis* (**Supplementary Figure 7**, pdb files are available in supplementary material). The most drastic difference between the *E.coli* and *C.maquilingensis* enzymes is the absence of cydB. cydB in *E.coli* was shown to contain the oxygen diffusion channel^31,32^ and an additional proton channel leading to heme *d*, bound to subunit I^30–32^. In *C. maquilingensis* the second subunit is cydA’ which is 26 % similar to cydA. Only the first two helices are well conserved between these subunits in *C. maquilingensis* whereas other qOR4a-type cydA’ and qOR4b-type cydA, such as in *A. fulgidus* are similar in the first 4 helices. To substitute for the proton channel that exists in cydB, conserved residues in cydA’ such as Thr71, Thr74 and His126 might form a different proton channel. cydA’ probably hosts an oxygen diffusion channel to substitute for the loss of the one in cydB but it is not possible to tell from the sequence alignment or structural model where in the subunit this might be. Interestingly, cydA’ retains His19 which has been implicated as a ligand to heme *d* and heme *b_595_* in *E. coli* and *G. thermodenitrificans* cytbd respectively, which might suggest that an additional heme might bind to the cydA’ subunit but we cannot verify or refute this from our protein preparation. A number of mutations are observed around the binuclear-active site in subunit I, which might affect the midpoint potential of the heme or the proton-coupled electron transfer mechanisms.

### Evolution of cydA’ and other cydA homologs missing the quinol binding site

As mentioned earlier, a phylogenomic analysis of cydA homologs revealed two new families, OR-C and OR-N that share the first four helices containing the oxygen reduction site. The cydA subunit of OR-C *bd*-type oxygen reductases typically has eight transmembrane helices and an extended C-terminal periplasmic portion that binds hemes *c*, strongly suggesting that a cytochrome *c* could be an electron donor to this family. Adjacent to the OR-C1a-type cydA is OR-C1b which also has 8 transmembrane helices. The OR-N3a/b, -N4a/b and -N5a/b family cydA typically have 10 helices while the OR-N2 and -N1 have 14 transmembrane helices. OR-N enzymes have been previously noted in Nitrospira^51^ and Chloroflexi (N5a/b)^52^. They were recently shown to be expressed in manganese oxidizing autotrophic microorganism, *Candidatus manganitrophus noduliformans* (N2/N1)from the phylum *Nitrospira^53^* and is implicated in oxygen reduction. Greater details on the OR-C and OR-N families, including distribution, alignments and conserved amino acids are found in **Supplementary Material**. The OR-C and OR-N families are widely distributed in Bacteria. OR-C is present only in Bacteria, while OR-N has very few representatives in Archaea (**Supplementary Table 2, Supplementary Table 4**).

A phylogenetic tree of all cydA clades suggest that the OR-C and OR-N families are more closely related to qOR4b than the other qOR subfamilies (**Figure 1**). There are also conserved sequence features that suggest that the OR-C and OR-N families are more closely related to the qOR3 and qOR4a families than the qOR1 family including the deletion in the proton channel between the conserved glutamates E101 and E108 like in *G.thermodenitrificans* cytbd, and the presence of nearby conserved tyrosines (Y123 and Y125 in the *G.thermodenitrificans* cytbd numbering). (**Supplementary table 6**). This suggests that the OR-C and OR-N families diverged from either of these two families and evolved after the qOR reductases. The evolutionary analysis within this family is complicated by the high number of independent gene duplication events. It appears that qOR4b, OR-N5b, OR-N3b, OR-N4b and OR-C1b subfamilies were the result of gene duplication events (**Figure 1**). In fact, OR-3a and OR-3b cydA share 50 % sequence similarity. Additionally, the presence of OR-N1-type and OR-N2-type cydA in the same operon in some Bacteria and the extent of similarity between them (up to 40%) suggest that they were part of yet another gene duplication. The importance of gene duplication in protein evolution and functional diversification is well-known^54^. The nature this process has taken in the cytochrome *bd*-type oxygen reductases is interesting - a majority of the cydA paralogs have maintained the architecture associated with oxygen reduction and all of them have maintained the His19 ligand to the active site heme *d* (as per the *E. coli* structure). Additionally, all the above-mentioned duplication events appear to have resulted in a complex of multiple cydA-like proteins with the possible exception of OR-N1 and OR-N2. OR-N2 is often found in operons without another cydA-like protein (**Figure 1**). His19 and heme *d* is found near the interface of subunit I and subunit II in the cydAB structures and the complete conservation of these features with a change in their interacting partner, is suggestive of the process of duplication and interface evolution recently investigated in hemoglobin^55^. Future work in the biochemical and structural characterization of the various cytbd families will help us develop insight into the driving forces behind the evolution of this superfamily. Presently, it is clear that the defining characteristic of the cytbd superfamily is the di-heme oxygen reduction site found in the first four helices of cydA homologs. Our analysis suggests that the *bd* protein scaffold was diversified multiple times to perform O_2_ chemistry in unique environments, possibly to function with different electron donors.

## Conclusions

The superfamily of cytochrome *bd*-type oxygen reductases is one of only two oxygen reductase superfamilies that are widely distributed in Bacteria and Archaea. In the current work we have demonstrated the large diversity of this superfamily using phylogenomics. In addition, we biochemically characterized the CydAA’ from *C. maquilingensis* showing that cydAA’ is a robust oxygen reductase. The isolated CydAA’ contained only *b* hemes and no heme *d*. Hence, *C. maquilingensis* CydAA’ is a *bb*’-type oxygen reductase and is the first such enzyme to be purified and demonstrated to be a functional oxygen reductase. Finally, we demonstrate that significant diversification of the cydA has occurred with the conserved oxygen reduction site being adapted to multiple functions within various ecological niches.

## Materials and Methods

### Phylogenomic analysis of cytochrome *bd* sequences in the GTDB database

In order to reconcile the protein phylogeny of cytochrome *bd* oxygen reductases with species taxonomy, we identified and mapped all cytbd to their respective species within the GTDB database release89^37^. All cydA sequences were extracted from GTDB genomes using BLAST^56^ with an e-value of 1e-1. The sequences were then aligned using muscle^57^ using the optional maxiters cut-off of 2. The alignment was visualized on Jalview^58^ and sequences were filtered to remove cydA sequences without characteristics of the quinol binding site or the proton channel. This filtration step was used to remove subunits II but also resulted in the loss of a few subunits I within qOR1 that appear to have lost the proton channel. The filtered set of cydA sequences were then classified using a Hidden Markov Model (HMM)-based classifier trained to identify the families – qOR1, qOR2, qOR3, qOR4a, OR-C and OR-N. The HMMs for those subfamilies and families are available in the supplementary material. The presence or absence of cytbd in each species was tabulated and is available as **Supplementary Table 2**. The all archaea species tree used to analyze the distribution of cytochrome *bd* oxygen reductases in archaea was generated using Anvi’o^41^. A multiple sequence alignment was created by extracting all ribosomal proteins from archaeal genomes using the HMM source Archaea_76. This alignment was then used to generate a phylogenetic tree using FastTree as per Anvi’o’s default settings. This tree was annotated using the data available in **Supplementary Table 4** on the iTOL server^59^.

The protein phylogeny of cytbd sequences was inferred using sequences of cytbd subunit I, cydA. These were extracted from a taxonomically diverse set of genomes and metagenomes from IMG^60^, filtered with UCLUST^61^ using a percentage identity cut-off of 0.6 and aligned using MUSCLE. The multiple sequence alignment was used to infer a phylogenetic tree using RAxML^62^ on the CIPRES Science Gateway^63^ with the PROTGAMMA substitution model, DAYHOFF matrix specification and a bootstrap analysis with 100 iterations.

### Preparation of construct for of cytochrome *bd* oxidase from *Escherichia coli*

The genes encoding the *bb*’ oxygen reductase (Gene Object ID: 641276193-4) from *C. maquilengensis* were PCR amplified using primers purchased from Integrated DNA Technology. The genes were cloned into pET22b (Invitrogen) using 5’ NdeI and 3’ XhoI cut sites. The inherent 6-Histidine tag in the vector was used to purify the protein. The vector was engineered to use EGFP, GST or FLAG tags alternatively. The tag was added to subunit II in case of EGFP and FLAG; a tag on both subunit I and II was attempted for the His-tag and GST tag. The expression vector, along with pRARE (Novagen) was then transformed into (CBO ΔcydBΔappC::kan) for protein expression.

### Cell Growth and Protein Purification

A single colony was inoculated into 5 ml of LB (yeast extract and tryptone were purchased from Acumedia and NaCl from Sigma-Aldrich) with 100 μg/ml Ampicillin and incubated with shaking at 37 °C. The following day, the 5 ml culture was inoculated in 300 ml LB with 100 μg/ml Ampicillin and grown overnight at 37 °C. On the third day, 10 ml of the secondary culture was inoculated into twenty four of 2L flasks containing 1 L LB with 100 μg/ml Ampicillin, each. The flasks were incubated at 37 °C while shaking at 200 rpm, until the OD600 of the culture reached 0.6. The temperature was then lowered to 30 °C, and the culture was incubated for 8 hrs or overnight.

The fully-grown cultures were then pelleted by spinning down at 8000 rpm for 8 minutes, in 500 ml centrifuge bottles. The harvested cells were then resuspended in 100 mM Tris-HCl, 10 mM MgS04, pH 8 with DNaseI and a protease inhibitor cocktail from Sigma. The cells were then homogenized using a Bamix Homogenizer, and passed through a Microfluidizer cell at 100 psi, three times, to lyse the cells. The soluble fraction of the lysate was then separated from the insoluble by spinning down the lysate at 8000 rpm. Membranes were extracted from the soluble fraction by centrifuging the soluble fraction at 42000 rpm for 4 hours.

Membranes were resuspended in 20 mM Tris, 300 mM NaCl, pH 8 and then solubilized with 1% DDM or 1% SML. The solubilized membranes were spun down at 42000 rpm for 45 minutes to remove unsolubilized membranes. The supernatant was stirred with Ni-NTA resin for 1 hr and then loaded onto a column. The flow through was shown to contain the cydAA’ because of its poor affinity for the nickel column. The flow through was then diluted in buffer to contain 50 mM salt and then loaded onto a DEAE column equilibriated with 20 mM Tris, pH 8, 0.05% DDM. An elution gradient was run between 0-500 mM NaCl and cydAA’ was partially purified from the fraction with higher absorbance at A_412nm_, corresponding to the soret peak for heme *b* and used for assays and spectroscopy. This is similar to the first step for purification of *Escherichia coli*.

### UV-visible spectroscopy

Spectra of the protein were obtained using an Agilent DW-2000 Spectrophotometer in the UV-visible region. The cuvette used has a pathlength of 1cm. The oxidized spectrum was taken of the air-oxidized protein. The enzyme was reduced with dithionite to obtain a reduced spectrum.

### Collection of Pyridine Hemochrome spectra and Heme Analysis

For the wildtype or mutants enzymes, 35 μl of the enzyme solution was mixed with an equal volume of 40% pyridine with 200 mM NaOH. The oxidized spectra was measured in the presence of ferricyanide and the reduced in the presence of dithionite. The values of heme *b* were calculated according to the matrix suggested in^46^. The concentration of heme *d* was estimated using the extinction coefficient ε_(629-670nm)_ = 25 mM^−1^cm^−1^.

### Measurement of oxygen reductase activity

Oxygen reductase activity was measured using the Mitocell Miniature Respirometer MT200A (Harvard Apparatus). 5 mM DTT and 350 μM Q1 were used as electron donors to measure oxygen reduction by *C.maquilingensis* cydAA’ and *E.coli* cytochrome *bd*. 150-250 μM KCN was used to test the cyanide sensitivity of the enzymes.

### Structural modelling of cydAA’ from *Caldivirga maquilingensis*

Sequences of subunit I from *Geobacillus thermodenitrificans* and *Caldivirga maquilingensis* were aligned using a larger alignment comprising many hundreds of *bb*’ sequences made with the software MUSCLE. This alignment was used as to create a model of subunit I from *Caldivirga* using the *Geobacillus* subunit I as a template on the Swiss Model server. A model of subunit II was also created using subunit I as a template. (The alignments are provided as Supplementary Figures S4 and S5) The model was then visualized using VMD 1.9.2beta1.

## Supporting information

Supplementary Table 1. Total number of various cydA families and subfamilies in the GTDB.

Supplementary Table 2. Distribution of cydA subfamilies by genome in GTDB.

Supplementary Table 6. Conserved features residues identified in cytbd families without quinol oxidizing features - qOR4b and subfamilies OR-C1a, OR-N

Supplementary Table 7. Number of heme c binding sites in cytbd sequences from OR-C1a/OR-C1b subfamilies.

Supplementary Table 3. Distribution of cydA subfamilies by phyla in GTDB.

Supplementary Table 4. Distribution of quinol oxidizing cytbd in archaeal genomes found in the GTDB.

## List of Tables

**Table 1. Expression of cydAA’ in the environment estimated using publicly available metatranscriptomes.** cydA homologs from the metatranscriptomic data available on the IMG website were extracted using a BLASTP cutoff of 1e-5. The short fragments found in metatranscriptomic data were matched with the full corresponding protein sequence based on the best hit in the NCBI database.

**Table 2. Oxygen reduction activity of *E. coli* cydAB and *C. maquilingensis* cydAA’ in the presence of 350 μM coenzyme Q1 and 5 mM DTT.**

**Supplementary Table 1. Total number of various cydA families and subfamilies in the GTDB**. All cydA sequences were extracted from GTDB genomes using BLAST with an e-value of 1e-1. The sequences were then filtered to remove cydA sequences without characteristics of the quinol binding site and then classified using a Hidden Markov Model (HMM)-based classifier trained to identify the families – qOR1, qOR2, qOR3, qOR4a, OR-C, OR-N. The total number of cydA sequences in each of these families and subfamilies were summed to generate this table.

**Supplementary Table 2. Distribution of cydA subfamilies by genome in GTDB**. All cydA sequences were extracted from GTDB genomes using BLAST with an e-value of 1e-1. The sequences were then filtered to remove cydA sequences without characteristics of the quinol binding site and then classified using a Hidden Markov Model (HMM)-based classifier trained to identify the families – qOR1, qOR2, qOR3, qOR4a, OR-C, OR-N. Labelled cydA sequences were then mapped back to each species to generate this table.

**Supplementary Table 3. Distribution of cydA subfamilies by phyla in GTDB**. All cydA sequences were extracted from GTDB genomes using BLAST with an e-value of 1e-1. The sequences were then filtered to remove cydA sequences without characteristics of the quinol binding site and then classified using a Hidden Markov Model (HMM)-based classifier trained to identify the families – qOR1, qOR2, qOR3, qOR4a, OR-C, OR-N. Labelled cydA sequences were then mapped back to each species and the cydA sequences from each family/subfamily were summed across each phyla.

**Supplementary Table 4. Distribution of quinol oxidizing cytbd in archaeal genomes found in the GTDB**. All cydA sequences were extracted from GTDB genomes using BLAST with an e-value of 1e-1. The sequences were then filtered to remove cydA sequences without characteristics of the quinol binding site and then classified using a Hidden Markov Model (HMM)-based classifier trained to identify the families – qOR1, qOR2, qOR3, qOR4a. Labelled cydA sequences were then mapped back to each species within the domain archaea to generate this table.

**Supplementary Table 5. Growth conditions and temperature for organisms containing cydAA’**. Growth conditions, sensitivity to oxygen and temperature are detailed with references in this table to see if there is a pattern to the conditions under which cydAA’ is typically found.

**Supplementary Table 6. Conserved features residues identified in cytbd families without quinol oxidizing features – qOR4b and subfamilies OR-C1a, OR-N1, OR-N2, OR-N3a, OR-N3b, OR-N4a, OR-N4b, OR-N5a and OR-N5b**. Conserved residues were identified using multiple sequence alignments of cydA sequences from the above families. The presence or absence of the conserved residues for the three hemes, proton channel, quinol binding site are marked.

**Supplementary Table 7. Number of heme *c* binding sites in cytbd sequences from OR-C1a/OR-C1b subfamilies**. Number of heme *c* binding sites were counted using a python script that identifies the number of CXXCH motifs in each protein sequence. The OR-C1a sequences from Desulfovibrionia appear to have the greatest number of heme *c* binding sites – up to 8.

## List of multiple sequence alignments

**Supplementary multiple sequences alignment MSA1.** Multiple sequence alignment of sequences from cydA subfamilies qOR1, qOR2, qOR3, qOR4a, qOR4b and families OR-C1a and OR-N. OR-N is not split into subfamilies in this alignment. Various families and subfamilies are grouped when visualized in Jalview and amino acids are colored using a ClustalX algorithim with a greater than 90 % identity. This alignment was manually curated to improve the alignment and reduce the number of gaps.

**Supplementary multiple sequences alignment MSA2.** Multiple sequence alignment of sequences from cydA subfamilies qOR1, qOR2, qOR3, qOR4a, qOR4b. OR-C1a, OR-N1, OR-N2. OR-N3a/3b, OR-N4a/4b, OR-N5a and OR-N5b. Various families and subfamilies are grouped when visualized in Jalview and amino acids are colored using a ClustalX algorithim with a greater than 90 % identity.

**Supplementary multiple sequences alignment MSA3.** Multiple sequence alignment of sequences from all 15 cydA subfamilies qOR1, qOR2, qOR3, qOR4a, qOR4b. OR-C1a, OR-C1b OR-N1, OR-N2. OR-N3a/3b, OR-N4a/4b, OR-N5a and OR-N5b.

## List of supplementary figures

**Supplementary Figure 1. Phylogenetic clustering of all cydA-like sequences.** cydA sequences were extracted from a taxonomically diverse set of genomes and metagenomes from IMG and aligned using MUSCLE. The multiple sequence alignment was used to infer a phylogenetic tree using RAxML. The RAxML tree topology was similar to that inferred by PhyML. The three families, qOR, OR-C and OR-N are clearly separated, and 15 subfamilies were designated based on the clustering observed and identifiable sequence characteristics.

**Supplementary Figure 2. Sequence characteristics of qOR4a-cydA.** a. An unrooted phylogenetic tree of cydA sequences from archaea including both qOR4a-type cydA and cydA of the qOR1, qOR2 and qOR3 (in red) types was generated using RaxML. qOR4a type cydA have some internal clusters, identified with a shaded box in green, blue and purple. Characteristics unique to the cluster, when identifiable were indicated. For e.g., the presence of the insertion in the proton channel. b. A multiple sequence alignment of the sequences present in the above clusters, the background of each sequence cluster shaded in red, green, blue and purple according to the colors in the tree in a. Conserved residues corresponding to the ligands, proton channel and a few residues expected to participate in proton-coupled electron transfer are marked. Significantly, several qOR4a-type cydA have a lysine substituted for M393 (*E. coli* numbering) in the active site suggesting an alteration of the midpoint potential of heme *b_558_* in those enzymes.

**Supplementary Figure 3. GFP-tagged cydAA’ from *Caldivirga maquilingensis*.** The presence of cydAA’ during protein purification protocol was verified by looking at elution fractions under UV-light. Three glass vials containing (from leftmost) elution buffer, an elution fraction containing GFP-tagged cydAA’ and a fraction without cydAA’ are compared. The green fluorescence in the cydAA’ containing fraction is easily distinguishable.

**Supplementary Figure 4. Mass spectrometric identification of subunit I of cydAA’ from *Caldivirga maquilingensis*.** Partially purified cydAA’ was digested with Chymotrypsin and the digested peptides were separated by HPLC and infused into a Thermo LTQ Velos ETD Pro Mass Spectrometer. The mass fragments recovered after MS/MS fragmentation were subject to analysis by Mascot Distiller and Mascot version 2.4. The analysis revealed peptides from subunit I of cydAA’ in the protein preparation with a MASCOT score of 85.

**Supplementary Figure 5. SDS-PAGE gel electrophoresis of partially purified cydAA’.** A. Cell lysate was loaded onto a Ni-NTA column (Lane 1). The flow-through was loaded onto a Q-sepharose column and subject to elution by changing the salt concentration from 0-500 mM NaCl. Three elution peaks (Lanes 2,3,4) which absorbed at 412 nm were pooled, concentrated and diluted to 50 mM NaCl and then loaded onto a DEAE-Sepharose column and subject to elution under a salt gradient from 0-500 mM NaCl. The elution fraction (Lane 6) which absorbed at 412 nm was pooled and concentrated and used to identify electrophoresis patterns. Assays and spectra were obtained with a sample that was subject to a simpler purification protocol – Ni-NTA followed by DEAE-sepharose because the yield was poor from the 3-step purification protocol. Lane 5 was the Precision Plus Dual Color Standard (Bio-Rad). Subunits I and II are similar in size to the subunits from *E.coli* cytbd (B). B. Electrophoresis patterns of purified *E.coli* cytbo_3_ and cytbd.

**Supplementary Figure 6. Sequence characteristics of qOR4b-cydA from *Caldivirga maquilingensis*** (also referred to as cydA’). a. A topological representation of cydA’ using HMMTOP. The amino acids conserved above 90% identity are shaded in black. b. A multiple sequence alignment of sequences from qOR4a-cydA and qOR4b-cydA family. The former sequences are highlighted with the purple background while the latter are highlighted with a gray background. The absence of the proton channel residues E99 and E109 is apparent. The ligand to heme *d*, His19 and the proton channel residue H126 are completely conserved. The ligands to heme *b_558_* (H186 and M393) and other amino acids typically associated with the quinol binding site in helices V-VIII are not well conserved. Two threonines Thr71 and Thr74 in *C.maquilingensis* which take the place of Leu71 and Glu74 are completely conserved.

**Supplementary Figure 7. Structural model of subunits I from *Geobacillus thermodenitrificans* and *Caldivirga maquilingensis* respectively.** The homology model of cydA from *Caldivirga maquilingensis* was generated using the Swiss PDB viewer and visualized using VMD. The boxed regions reveal more polar residues in *Geobacillus*, represented by red and blue, while aromatic residues are colored in green.

