## Supplementary Material for "Evolution of the cytochrome-*bd* type oxygen reductase superfamily and the function of cydAA’ in Archaea"

#### Description of the various families and subfamilies within the cytochrome *bd*-type oxygen reductases

The cytochrome *bd* oxygen reductase superfamily is divided into 3 families based on phylogenetics and structure – qOR, OR-C and OR-N. qOR is defined by the presence of the quinol binding site in subunit I (*cydA*). OR-C is missing the quinol binding site but has a heme *c* binding site in subunit I. All of the OR-C family cytb<sub>d</sub> have heme *c* binding sites, identified by the presence of the CXXCH motif. The electron donor to the OR-C family is likely cytochrome *c* as is indicated by the presence of the heme *c* binding site. Enzymes from these family are widely distributed in the phyla Acidobacteria, Desulfobacterota, Desulfobacterota\_A, Campylobacterota and Desulfuromonadota. The number of heme *c* binding sites varies within the various cytb families and subfamilies - OR-C1a from Desulfovibrionia especially appear to have a great many heme *c* binding sites, as many as 8 (Supplementary Table S7). The presence of many hemes *c* could be typical of many cytoplasmic membrane complexes in deltaproteobacteria<sup>1</sup>. The ligand to heme *d*/heme *b*<sub>595</sub>, H19 is conserved and so are the glutamates E99 and E107 (*E.coli* cytb numbering) but of H126 and S140, other residues typically found in the proton channel, the former is not well conserved while the latter is sometimes a threonine.

OR-N is also missing the quinol binding site and is commonly found in operons containing alternative electron donors. At least 9 subfamilies the OR-N family are also shown (Supplementary Figure 1). The operon context and putative protein complex arrangement of each cydA-containing enzyme is also shown with a reference protein accession number and source microorganism. While OR-N3a and OR-N3b and OR-N4a and OR-N4b are found by themselves in the genome, OR-N1, OR-N5a/5b and OR-N2 are found with several other subunits within the same operon. OR-N1 and OR-N5a/b are typically found with petA, petB and petD subunits. These subunits are homologous respectively to the domains in cytochrome bc<sub>1</sub> complex and cytochrome b<sub>6</sub>f complex. petA contains the Fe/S cluster, while petB typically contains heme *b*. petD has been implicated in binding *Chl*a in the spinach b<sub>6</sub>f complex<sup>2</sup>. This would suggest that at least OR-N1 and OR-N5a/b complexes use quinol as an electron donor like cytochrome bc<sub>1</sub> and cytochrome b<sub>6</sub>f complex. OR-N2 cydA containing operons typically multiple cytochromes *c* and a subunit homologous to ebdC, the gamma subunit of ethyl benzene dehydrogenase which is implicated as a potential membrane anchor for the dehydrogenase which contains an Mo-Fe-S co-factor. It is not clear what the electron donor to this enzyme is but the presence of heme c binding subunits might suggest that it is cytochrome c. There is significant variation amongst the residues conserved within the proton channel for each of the OR-N subfamilies (Supplementary Table 6). However, some proton channel residues and the H19 (*E.c* numbering) ligand to active site heme are present like most *cydA* sequences from various *cytbd* subfamilies and families.

#### **Evolutionary analysis of structural differences between the enzymes from the subfamilies qOR1 and qOR3**

A vast majority of the *bd*-family diversity has not yet been biochemically characterized. However, what has been previously demonstrated does pose an interesting quandary. Previous structural evidence<sup>3-5</sup> has suggested that the active sites in different cytochrome *bd*-type oxygen reductases can have distinct active site heme arrangements. The location of heme *d* in *G. thermodenitrificans* of the qOR3-family is adjacent to Glu445 while heme *b*<sub>595</sub> is adjacent to His19<sup>3</sup>. In *E. coli* *cytbd* from the qOR1-family, these positions are reversed<sup>4,5</sup>. The presence of the fourth subunit *cydY/cydH/ynhF* and the insertion of Leu101 have been implicated in the differences between these two isoforms. It is not certain whether the presence of the fourth subunit is universal amongst qOR1 family of *cytbd* but, the insertion between the two conserved glutamates E99 and E107 in the qOR1 family is completely conserved. The EM structures of *cytbd* would seem to suggest this insertion leads to a curvature within that helix but the amino acid that is inserted varies a great deal – it is typically either Leu, Met, Thr, Val or Ile. The evolutionary distance between the qOR3 and qOR1 families is certainly consistent with the observation of biochemical differences between these characterized enzymes but it is not clear that any of these differences can be used to explain the differences in the placement of hemes, especially given previous evidence that the CIO-type of cytb<sub>d</sub> can be found with both hemes *b* and *d*<sup>6</sup> and that heme *d* is expected to be formed in-situ<sup>7</sup>.

#### **Conservation of Q-loop within the qOR1 subfamily**

It has been noted that the qOR1-family cytb<sub>d</sub> has a hydrophilic loop between transmembrane helices VI and VII, called the Q-loop, which is implicated in quinol oxidation<sup>8</sup>, while a shorter Q-loop is found in the qOR3-family enzyme from *Geobacillus thermodenitrificans*. This loop appears to contribute to quinol oxidation without directly affecting substrate binding<sup>4,5</sup>. Significantly, all the long Q-loop containing cytochrome *bd* belong to the qOR1 clade. In our observations, long Q-loop members do not appear to form a monophyletic clade of enzymes, suggesting that its insertion and deletion appear to have happened multiple times within this clade, which is interesting given its demonstrated importance for enzyme stability<sup>9</sup>. Since small Q-loop and long Q-loop cytochrome *bd* enzymes within the qOR1 clade can be evolutionarily closely related, the physiological implication of the presence of the loop is unclear. Structural characterization of short Q-loop enzymes from the qOR1 clade and other clades identified here would help in understanding the biochemical variability that comes from evolutionary diversification.

Cytochrome *bd*-type
oxygen reductase
superfamily

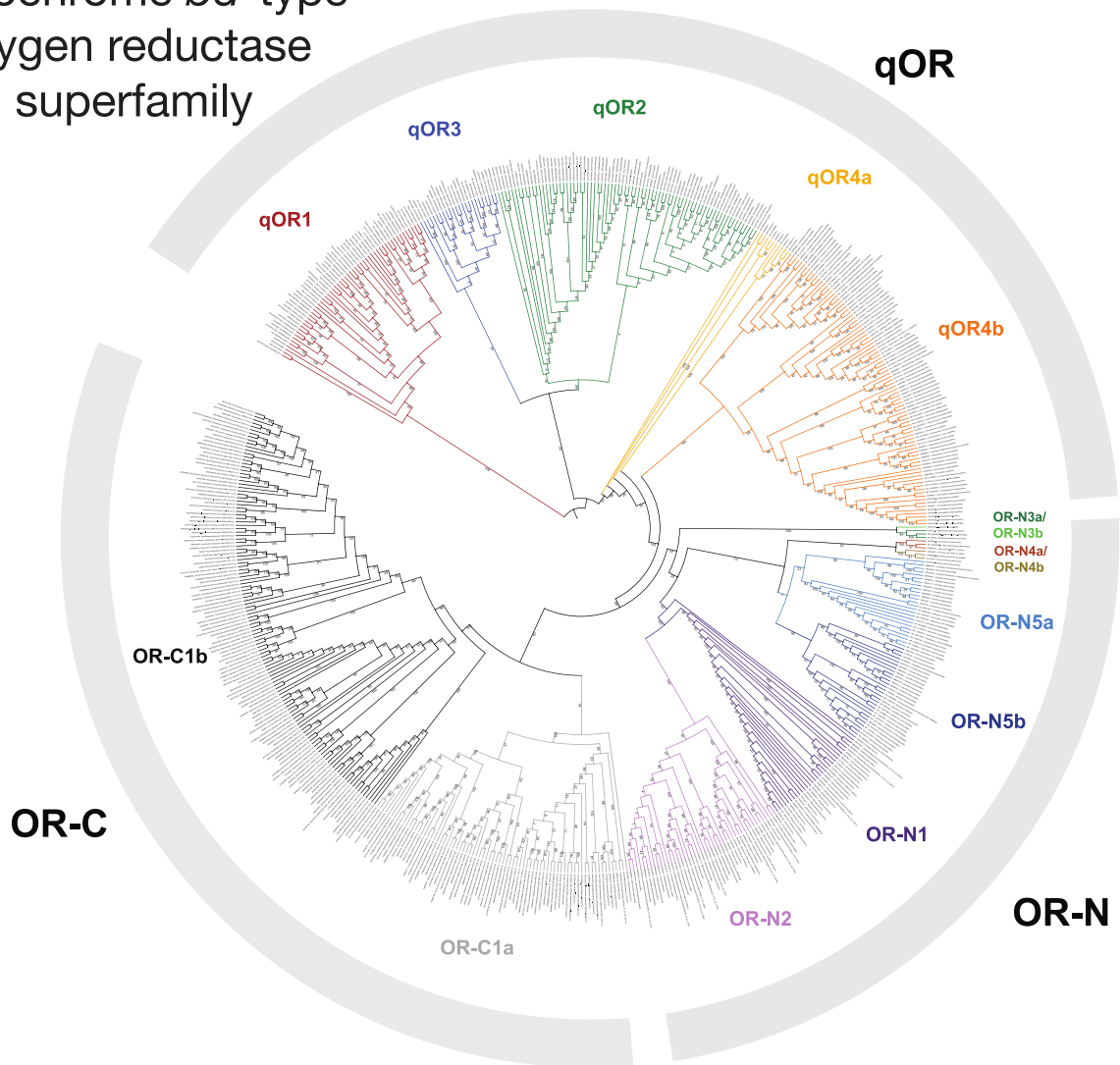

**Supplementary Figure 1. Phylogenetic clustering of all *cydA*-like sequences.** *cydA* sequences were extracted from a taxonomically diverse set of genomes and metagenomes from IMG and aligned using MUSCLE. The multiple sequence alignment was used to infer a phylogenetic tree using RAxML. The RAxML tree topology was similar to that inferred by PhyML. The three families, qOR, OR-C and OR-N are clearly separated, and 15 subfamilies were designated based on the clustering observed and identifiable sequence characteristics.

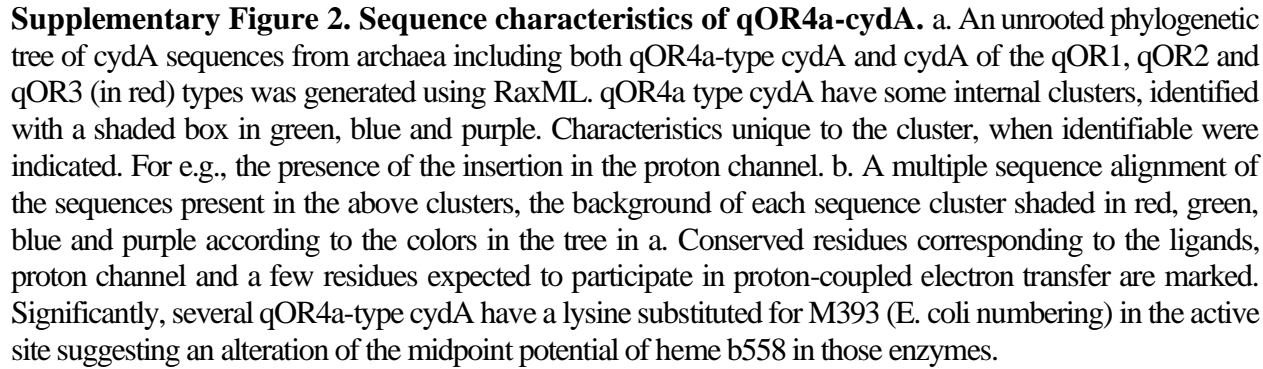

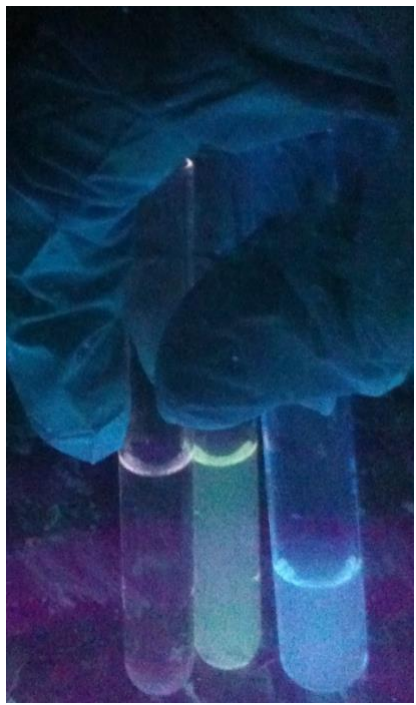

**Supplementary Figure 3. GFP-tagged cydAA' from *Caldivirga maquilingensis*.** The presence of cydAA' during protein purification protocol was verified by looking at elution fractions under UV-light. Three glass vials containing (from leftmost) elution buffer, an elution fraction containing GFP-tagged cydAA' and a fraction without cydAA' are compared. The green fluorescence in the cydAA' containing fraction is easily distinguishable.

### Protein sequence coverage: 12%

Matched peptides shown in **bold red**.

```
1  MPSAYVLSII NASTLDAARW  VSAIGILAH  LSLASSFLGTI  LIAVIAEYLY
51  LVKHKDKDWYD  KARMFSVVST  IFFGVGAAG  TLVEFGLVTI  WSNFITIIGS
101 AIVLPFYLEL  FAFLTEVILL  PLYVFTWGV  RNGVWHWVIG  LAAAFGGYWS
151 AYNILAVMAS  LSMRPPGMII  QNLAASNETI  AGLTSYLVTV  AKPTDAWNMF
201 WWGANVIFH GILAAAILTW  SVVGAIYLYG  YVREHRPDQA  KVLKVIIPGV
251 AVMTAIEGFI  LGHDQGELVL  QFDPLKLA  EGMFWKGLKV  DPLLSFFAYG
301 TFNHAFWGY  SWPANVRPPL APFDFLPIFY  LGFMVTLGVL  LGVWSGGLSL
351 WYLFNGFFSR  FNWARVIASF  LERTGPYAMP  LFAALAAIGG AVTSESGRVP
401 FILVDSSNP  NGGPPTVTGV  PIYEGGLINP  NLSLPGWLVA  LIIIVEVAMP
451 ALAVYMVYLY  TKPKEVKPTQ  VVEY
```

**Supplementary Figure 4. Mass spectrometric identification of subunit I of cydAA' from *Caldivirga maquilensis*.** Partially purified cydAA' was digested with Chymotrypsin and the digested peptides were separated by HPLC and infused into a Thermo LTQ Velos ETD Pro Mass Spectrometer. The mass fragments recovered after MS/MS fragmentation were subject to analysis by Mascot Distiller and Mascot version 2.4. The analysis revealed peptides from subunit I of cydAA' in the protein preparation with a MASCOT score of 85.

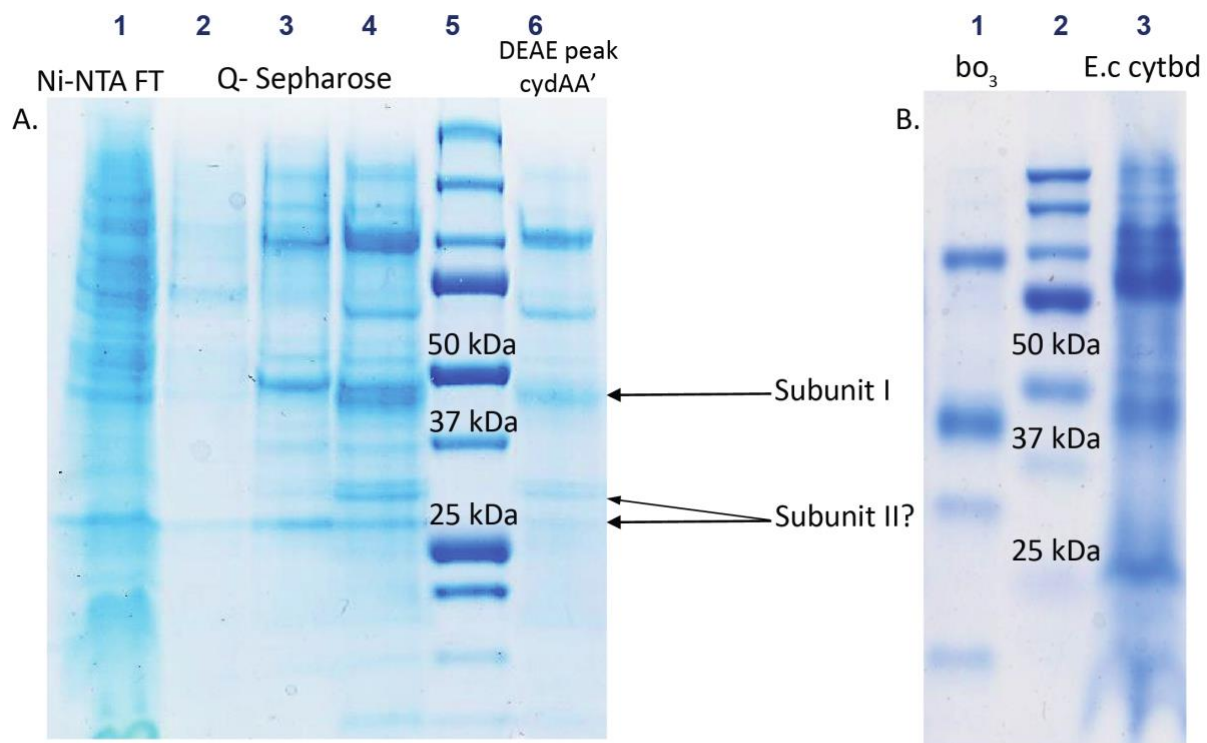

**Supplementary Figure S5. SDS-PAGE gel electrophoresis of partially purified *cydAA'*.** A. Cell lysate was loaded onto a Ni-NTA column (Lane 1). The flow-through was loaded onto a Q-sepharose column and subject to elution by changing the salt concentration from 0-500 mM NaCl. Three elution peaks (Lanes 2,3,4) which absorbed at 412 nm were pooled, concentrated and diluted to 50 mM NaCl and then loaded onto a DEAE-Sepharose column and subject to elution under a salt gradient from 0-500 mM NaCl. The elution fraction (Lane 6) which absorbed at 412 nm was pooled and concentrated and used to identify electrophoresis patterns. Assays and spectra were obtained with a sample that was subject to a simpler purification protocol – Ni-NTA followed by DEAE-sepharose because the yield was poor from the 3-step purification protocol. Lane 5 was the Precision Plus Dual Color Standard (Bio-Rad). Subunits I and II are similar in size to the subunits from *E.coli* *cytb3* and *cytbD*. B. Electrophoresis patterns of purified *E.coli* *cytbo3* and *cytbD*.

a. A representation of the transmembrane helices in *cydA'* generated by HMMTOP.

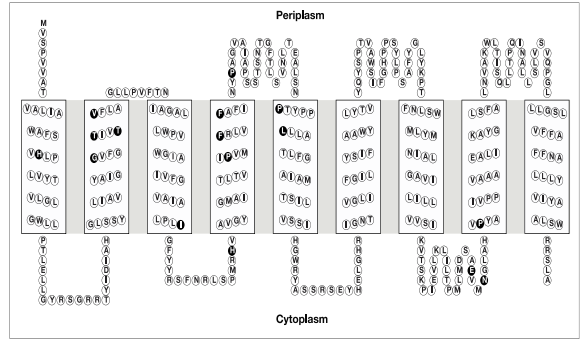

b. Sequence alignment of *cydA'* and *cydA* (including *cydA*)

|  | **E. coli *cydA* numbering** | **H19** |  | **H186** |
| --- | --- | --- | --- | --- |
| Thermoproteus tenax | -----MPG-----PLD-----LELARLSAVG-----VLAMGLASSFLGALGV |  | Thermoproteus tenax | ...QIVNLA-----AGKSIAGAT-VLATTC-NPADANNIFWGANVFIFGILL |
| Volcaniseta sp. EB80 | -----MTD-----QILGAPTVLVITATQ-----VLAMGLASSFLGALGV |  | Volcaniseta sp. EB80 | ...EVINILY-----QATGQNVVGLTD-VVTKA-SPADANNIFWGANVFIFGILL |
| Volcaniseta moutnovskia | -----MDQPLGA-----PTD-----TRVSAVG-----ILAMGLASSFLGTLIAI |  | Volcaniseta moutnovskia | ...EVINILY-----QATGQNVVGLTD-VVTKA-SPADANNIFWGANVFIFGILL |
| Caldivirga maquilingensis | -----MFSAVLSINAS-----PLD-----IAKRVSAIG-----ILAMGLASSFLGTLIAI |  | Caldivirga maquilingensis | ...LQNLA-----ASNETIAGLTS-VLVTKA-KPTDANNIFWGANVFIFGILL |
| Pyrobaculum sp. WP30 | MTLSTGGGGLVLYSHYTH-----SAI-----IALAISISFQ-----TILRVYVGLIAITLIP |  | Pyrobaculum sp. WP30 | ...TLGAGSALISVNFQSVGFPAQ-NHLLA-NP-----AI-----STVWELL |
| Archaeoglobus fulgidus DSM 4304 | -----MGSG-----TII-----DAMLLGLSALYIINAVLITIGLPIVII |  | Archaeoglobus fulgidus DSM 4304 | ...PI-----QAIQNFVAKGVGLTD-VYALP-NP-----AA-----ISALRDL |
| Caldisphaera sp. | -----NVL-----VSFVLAFN-----FGHIVLVNLVIGLIALVF |  | Caldisphaera sp. | ...EFS-----CQ-----NHPPV-NVAMATNP-----PI-----PLYKSTV |
| Archaeoglobus sulfatocalidus | -----HIL-----YFIALV-----FGHIVLVNLVIGLIALVF |  | Archaeoglobus sulfatocalidus | ...ILT-----PKPHLD-VIAAFI-NP-----PI-----PLYKSTV |
| Pyrodicticum occultum | -----HLGRVAMERGDGV-----TAPAFALN-----FGHIVLVNLVIGLIALVF |  | Pyrodicticum occultum | ...GYDFVR-----RHEPLD-VGALITNP-----PI-----PLYKSTV |
| Candidatus Methanodesulfokores washburnensis | -----HIL-----YFIALV-----FGHIVLVNLVIGLIALVF |  | Candidatus Methanodesulfokores washburnensis | ...PI-----RHEPLD-VGALITNP-----PI-----PLYKSTV |
| Thermofilum sp. N113 | -----MEA-----PVP-----FLALV-----FGHIVLVNLVIGLIALVF |  | Thermofilum sp. N113 | ...SFD-----GKFWLD-VAKALA-NP-----PI-----PLYKSTV |
| Thermocladium sp. ECH B | -----N-----AAT-----TAVALLMA-----FSHIVLVNLVIGLIALVF |  | Thermocladium sp. ECH B | ...QITSS-----SSMIGELNVLGQALG-NP-----PI-----PLLIATLA |
| Thermoproteus sp. JCHS 4 | -----HDL-----TA-----TAVALLMA-----FSHIVLVNLVIGLIALVF |  | Thermoproteus sp. JCHS 4 | ...VTAFD-----NSIVG-FDLGVQ-LWQALG-NP-----PI-----PLLIATLA |
| Volcaniseta moutnovskia | -----HDL-----TA-----TAVALLMA-----FSHIVLVNLVIGLIALVF |  | Volcaniseta moutnovskia | ...EYFES-----TSIVPGLI-NHGFALN-----PI-----PLLIATLA |
| Caldivirga maquilingensis | -----MVS-----PVV-----ATVALIMA-----FSHIVLVNLVIGLIALVF |  | Caldivirga maquilingensis | ...AIAPSS-----TSNTG-FPLSVN-VTALIS-NP-----PI-----PLLIATLA |
| Volcaniseta sp. EB80 | -----N-----IDL-----TLFSLGFT-----LHMLVFWNLVIGLIALVF |  | Volcaniseta sp. EB80 | ...DGDPS-----TSNTG-VTALIS-NP-----PI-----ALALMTM |
| Pyrobaculum sp. WP30 | -----N-----SAY-----LSFVLGFT-----LHMLVFWNLVIGLIALVF |  | Pyrobaculum sp. WP30 | ...EYFES-----TSIVPGLI-NHGFALN-----PI-----PLLIATLA |
| Archaeoglobus fulgidus DSM 8774 | -----HIS-----VGFPLGFA-----VLAMGLASSFLGALGV |  | Archaeoglobus fulgidus DSM 8774 | ...EYFES-----TSIVPGLI-NHGFALN-----PI-----PLLIATLA |
|  | **L71 E74** |  |  |  |
| Thermoproteus tenax | IAEYLMARRDEWNLAKRNPITATIPFGGAAPFTLVEGLVTINSFIAITGATL |  | Thermoproteus tenax | -----EJ-----PQVYIFILGMVAGILLGLTG |
| Volcaniseta sp. EB80 | VAEYLMARRDEWNLAKRNPITATIPFGGAAPFTLVEGLVTINSFIAITGATL |  | Volcaniseta sp. EB80 | -----SVR-----PQLVYFVFMVIFPGLLVWGA |
| Volcaniseta moutnovskia | VAEYLYFRKNOFYNTARTFSVITIPFGGAAPFTLVEGLVTINSFIAITGATL |  | Volcaniseta moutnovskia | -----EJ-----PQVYIFVFMVIFPGLLVWGA |
| Caldivirga maquilingensis | IAEYLYLVKDGKDYKARMSVSTIPFGGAAPFTLVEGLVTINSFIAITGATL |  | Caldivirga maquilingensis | -----NVRPLAPDFPIFYILGFMVAGILLGLTG |
| Pyrobaculum sp. WP30 | ANLVKMRGTGGDLYKARTLZAWAVNFAVGVTOTVVEGLLEIWPISILLSSGF |  | Pyrobaculum sp. WP30 | -----NCRSAVASLEP-----LAPVSAVITMVGSGILLAVA-AL |
| Archaeoglobus fulgidus DSM 4304 | GLLAKHSGSGDEFFAKIMPAVLINFAAGVGTLLVEGLGAMGGLTIAISAPA |  | Archaeoglobus fulgidus DSM 4304 | ARQGVGVVSEELQAMNTELAGICLACAKASRVAIVAAVIT-KIARGVIGVGS-AL |
| Caldisphaera sp. | LFETLLGRMSDVLGRKRMVYVYVGVGTATVFLFSLPFTFDVIGLGLM |  | Caldisphaera sp. | -----SVL-----PQVYIFVFMVIFPGLLVWGA |
| Archaeoglobus sulfatocalidus | LYLRKRGKSNYAYIKSLQMLNFAATYALGVGTATVFLFSLPFTFDVIGLGLM |  | Archaeoglobus sulfatocalidus | -----SVL-----PQVYIFVFMVIFPGLLVWGA |
| Pyrodicticum occultum | LYLRKRGKSNYAYIKSLQMLNFAATYALGVGTATVFLFSLPFTFDVIGLGLM |  | Pyrodicticum occultum | -----SVL-----PQVYIFVFMVIFPGLLVWGA |
| Candidatus Methanodesulfokores washburnensis | LYLRKRGKSNYAYIKSLQMLNFAATYALGVGTATVFLFSLPFTFDVIGLGLM |  | Candidatus Methanodesulfokores washburnensis | -----SVL-----PQVYIFVFMVIFPGLLVWGA |
| Thermofilum sp. N113 | LYLRKRGKSNYAYIKSLQMLNFAATYALGVGTATVFLFSLPFTFDVIGLGLM |  | Thermofilum sp. N113 | -----SVL-----PQVYIFVFMVIFPGLLVWGA |
| Thermocladium sp. ECH B | LYLRKRGKSNYAYIKSLQMLNFAATYALGVGTATVFLFSLPFTFDVIGLGLM |  | Thermocladium sp. ECH B | -----SVL-----PQVYIFVFMVIFPGLLVWGA |
| Thermoproteus sp. JCHS 4 | LYLRKRGKSNYAYIKSLQMLNFAATYALGVGTATVFLFSLPFTFDVIGLGLM |  | Thermoproteus sp. JCHS 4 | -----SVL-----PQVYIFVFMVIFPGLLVWGA |
| Volcaniseta moutnovskia | LYLRKRGKSNYAYIKSLQMLNFAATYALGVGTATVFLFSLPFTFDVIGLGLM |  | Volcaniseta moutnovskia | -----SVL-----PQVYIFVFMVIFPGLLVWGA |
| Caldivirga maquilingensis | LYLRKRGKSNYAYIKSLQMLNFAATYALGVGTATVFLFSLPFTFDVIGLGLM |  | Caldivirga maquilingensis | -----SVL-----PQVYIFVFMVIFPGLLVWGA |
| Volcaniseta sp. EB80 | LYLRKRGKSNYAYIKSLQMLNFAATYALGVGTATVFLFSLPFTFDVIGLGLM |  | Volcaniseta sp. EB80 | -----SVL-----PQVYIFVFMVIFPGLLVWGA |
| Pyrobaculum sp. WP30 | LYLRKRGKSNYAYIKSLQMLNFAATYALGVGTATVFLFSLPFTFDVIGLGLM |  | Pyrobaculum sp. WP30 | -----SVL-----PQVYIFVFMVIFPGLLVWGA |
| Archaeoglobus fulgidus DSM 8774 | LYLRKRGKSNYAYIKSLQMLNFAATYALGVGTATVFLFSLPFTFDVIGLGLM |  | Archaeoglobus fulgidus DSM 8774 | -----SVL-----PQVYIFVFMVIFPGLLVWGA |
|  | **E99 E107 H126** |  |  | **E445 R448** |
| Thermoproteus tenax | PPFLLE-PAFLMEVFLPLVPTWGRINPWLWFGIGLAAPG-----GYSAYNILAVMS |  | Thermoproteus tenax | ALASTGGAV-SAEGRYPFLVQVSTGTC-PPMI-----TGVP----- |
| Volcaniseta sp. EB80 | PPFLLE-PAFLMEVFLPLVPTWGRINPWLWFGIGLAAPG-----GYSAYNILAVMS |  | Volcaniseta sp. EB80 | AFAAIGGSV-SAEGRYPFLVQVSTGTC-PPMI-----TGVP----- |
| Volcaniseta moutnovskia | PPFLLE-PAFLMEVFLPLVPTWGRINPWLWFGIGLAAPG-----GYSAYNILAVMS |  | Volcaniseta moutnovskia | AFASIGAT-SAEGRYPFLVQVSTGTC-PPMI-----TGVP----- |
| Caldivirga maquilingensis | PPFLLE-PAFLMEVFLPLVPTWGRINPWLWFGIGLAAPG-----GYSAYNILAVMS |  | Caldivirga maquilingensis | AFASIGAT-SAEGRYPFLVQVSTGTC-PPMI-----TGVP----- |
| Pyrobaculum sp. WP30 | PPFLLE-PAFLMEVFLPLVPTWGRINPWLWFGIGLAAPG-----GYSAYNILAVMS |  | Pyrobaculum sp. WP30 | AVAAVAGM-AEAGR-----PMTV-----YGL----- |
| Archaeoglobus fulgidus DSM 4304 | PPFLLE-PAFLMEVFLPLVPTWGRINPWLWFGIGLAAPG-----GYSAYNILAVMS |  | Archaeoglobus fulgidus DSM 4304 | AVPSVLGM-VREVR-----PMTV-----YGL----- |
| Caldisphaera sp. | PPFLLE-PAFLMEVFLPLVPTWGRINPWLWFGIGLAAPG-----GYSAYNILAVMS |  | Caldisphaera sp. | IESMEN-----LAEFQI-----PMTV-----YGL----- |
| Archaeoglobus sulfatocalidus | PPFLLE-PAFLMEVFLPLVPTWGRINPWLWFGIGLAAPG-----GYSAYNILAVMS |  | Archaeoglobus sulfatocalidus | IESMEN-----LAEFQI-----PMTV-----YGL----- |
| Pyrodicticum occultum | PPFLLE-PAFLMEVFLPLVPTWGRINPWLWFGIGLAAPG-----GYSAYNILAVMS |  | Pyrodicticum occultum | IESMEN-----LAEFQI-----PMTV-----YGL----- |
| Candidatus Methanodesulfokores washburnensis | PPFLLE-PAFLMEVFLPLVPTWGRINPWLWFGIGLAAPG-----GYSAYNILAVMS |  | Candidatus Methanodesulfokores washburnensis | IESMEN-----LAEFQI-----PMTV-----YGL----- |
| Thermofilum sp. N113 | PPFLLE-PAFLMEVFLPLVPTWGRINPWLWFGIGLAAPG-----GYSAYNILAVMS |  | Thermofilum sp. N113 | IESMEN-----LAEFQI-----PMTV-----YGL----- |
| Thermocladium sp. ECH B | PPFLLE-PAFLMEVFLPLVPTWGRINPWLWFGIGLAAPG-----GYSAYNILAVMS |  | Thermocladium sp. ECH B | IESMEN-----LAEFQI-----PMTV-----YGL----- |
| Thermoproteus sp. JCHS 4 | PPFLLE-PAFLMEVFLPLVPTWGRINPWLWFGIGLAAPG-----GYSAYNILAVMS |  | Thermoproteus sp. JCHS 4 | IESMEN-----LAEFQI-----PMTV-----YGL----- |
| Volcaniseta moutnovskia | PPFLLE-PAFLMEVFLPLVPTWGRINPWLWFGIGLAAPG-----GYSAYNILAVMS |  | Volcaniseta moutnovskia | IESMEN-----LAEFQI-----PMTV-----YGL----- |
| Caldivirga maquilingensis | PPFLLE-PAFLMEVFLPLVPTWGRINPWLWFGIGLAAPG-----GYSAYNILAVMS |  | Caldivirga maquilingensis | IESMEN-----LAEFQI-----PMTV-----YGL----- |
| Volcaniseta sp. EB80 | PPFLLE-PAFLMEVFLPLVPTWGRINPWLWFGIGLAAPG-----GYSAYNILAVMS |  | Volcaniseta sp. EB80 | IESMEN-----LAEFQI-----PMTV-----YGL----- |
| Pyrobaculum sp. WP30 | PPFLLE-PAFLMEVFLPLVPTWGRINPWLWFGIGLAAPG-----GYSAYNILAVMS |  | Pyrobaculum sp. WP30 | IESMEN-----LAEFQI-----PMTV-----YGL----- |
| Archaeoglobus fulgidus DSM 8774 | PPFLLE-PAFLMEVFLPLVPTWGRINPWLWFGIGLAAPG-----GYSAYNILAVMS |  | Archaeoglobus fulgidus DSM 8774 | IESMEN-----LAEFQI-----PMTV-----YGL----- |

**Supplementary Figure 6. Sequence characteristics of qOR4b-*cydA* from *Caldivirga maquilingensis* (also referred to as *cydA'*).** a. A topological representation of *cydA'* using HMMTOP. The amino acids conserved above 90% identity are shaded in black. b. A multiple sequence alignment of sequences from qOR4a-*cydA* and qOR4b-*cydA* family. The former sequences are highlighted with the purple background while the latter are highlighted with a gray background. The absence of the proton channel residues E99 and E109 is apparent. The ligand to heme d, His19 and the proton channel residue H126 are completely conserved. The ligands to heme b558 (H186 and M393) and other amino acids typically associated with the quinol binding site in helices V-VIII are not well conserved. Two threonines Thr71 and Thr74 in *C.maquilingensis* which take the place of Leu71 and Glu74 are completely conserved.

**a.**

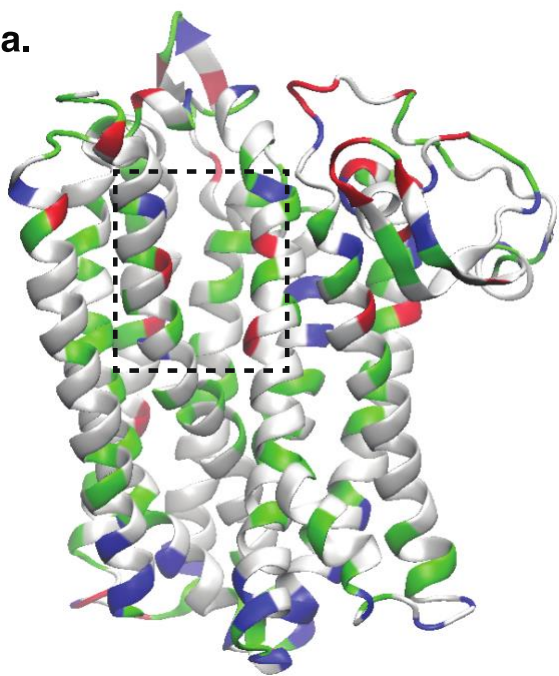

***Geobacillus thermodenitrificans***

**b.**

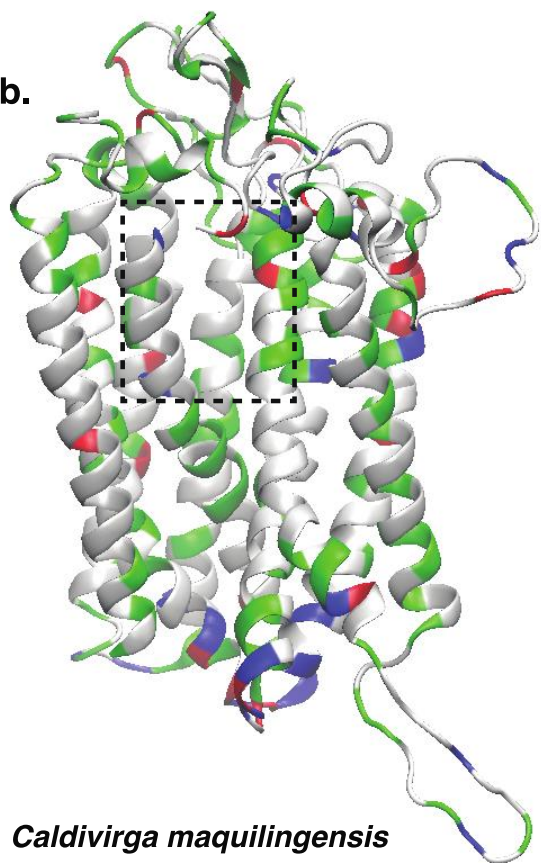

***Caldivirga maquilingensis***

**Supplementary Figure S7. Structural model of subunits I from *Geobacillus thermodenitrificans* and *Caldivirga maquilingensis* respectively.** The homology model of cydA from *Caldivirga maquilingensis* was generated using the Swiss PDB viewer and visualized using VMD. The boxed regions reveal more polar residues in *Geobacillus*, represented by red and blue, while aromatic residues are colored in green.
